# Stress relaxing granular bioprinting materials enable complex and uniform organoid self-organization

**DOI:** 10.1101/2024.02.01.578324

**Authors:** Austin J. Graham, Michelle W.L. Khoo, Vasudha Srivastava, Sara Viragova, Honesty Kim, Kavita Parekh, Kelsey M. Hennick, Malia Bird, Nadine Goldhammer, Jie Zeng Yu, Grace Hu, Natasha T. Brinkley, Lucas Pardo, Jasmine S. Amaya, Cameron D. Morley, Nishant Chadha, Paul Lebel, Sanjay Kumar, Jennifer M. Rosenbluth, Tomasz J. Nowakowski, Ovijit Chaudhuri, Ophir Klein, Rafael Gómez-Sjöberg, Zev J. Gartner

## Abstract

Complex and robust self-organization requires defined initial conditions and dynamic boundaries – neighboring tissues and extracellular matrix (ECM) that actively evolve to guide morphogenesis. A major challenge in tissue engineering is identifying material properties that mimic dynamic tissue boundaries but that are compatible with the engineering tools necessary for controlling the initial conditions of culture. Here we describe a highly tunable granular biomaterial, MAGIC matrix, that supports long-term bioprinting and gold-standard tissue self-organization. MAGIC matrix is designed for two temperature regimes: at 4 °C it exhibits reversible yield-stress behavior to support hours-long high-fidelity 3D printing without compromising cell viability; when transferred to cell culture at 37 °C, the material cross-links and exhibits viscoelasticity and stress relaxation that can be tuned to match numerous conditions, including that of reconstituted basement membrane matrices like Matrigel. We demonstrate that the timescale of stress relaxation and loss tangent are decoupled in MAGIC matrices, allowing us to test the role of stress relaxation rate and strain-dependence across formulations with identical storage and loss moduli. We find that fast absolute stress relaxation rates and large relative deformation magnitudes are required to optimize for morphogenesis. We apply optimized MAGIC matrices toward precise extrusion bioprinting of saturated cell suspensions directly into 3D culture. The ability to carefully control initial conditions for tissue growth yields dramatic increases in organoid reproducibility and complexity across multiple tissue types. We also fabricate perfusable 3D microphysiological systems that experience large strains in response to pressurization due to the compliant and dynamic tissue boundaries. Combined, our results both identify key parameters for optimal organoid morphogenesis in an engineered material and lay the foundation for fabricating more complex and reproducible tissue morphologies by canalizing their self-organization in both space and time.

## Introduction

Controlling the uniformity and complexity of tissues grown in vitro remains a major challenge. For example, organoids derived from the intestine lack key structural features found in vivo such as extended tubular geometries with continuous lumens, and manifest significant tissue-to-tissue heterogeneity^1–3^. Tissue in vivo typically self-organizes juxtaposed to other tissues that act as living boundaries, changing shape and composition in parallel to direct morphogenesis^4–8^. Guiding tissue self-organization in vitro requires replicating the important functional characteristics of these living interfaces, and satisfying these requirements remains a major challenge in materials science and tissue engineering^9,10^. Non-living biomaterials attempt to replicate key chemical and rheological cues derived from living boundaries, with gold-standards including reconstituted basement membrane matrices (rBMs) such as Matrigel as well as collagen I gels^11–13^. In the case of rBMs, recent studies have highlighted the presence of laminin epitopes as essential for setting epithelial polarity, with rheological properties such as low storage modulus and viscoelasticity being critical for morphogenesis^14–19^. This has enabled the design of synthetic biomaterials that support symmetry breaking during self-organization^1,20–24^. However, the morphology of tissues cultured in engineered materials has not been carefully compared to materials like Matrigel. Thus, the quantitative relationship between biomechanical properties like storage modulus, loss modulus, and stress relaxation on short and long timescales and the outcomes of morphogenesis remains unclear^25^.

An additional challenge to directing the uniform and complex self-organization of tissues in vitro is setting the initial conditions of a culture — defined here as the total number of cells, the proportions of each cell type, and their rough positions in 3D space. In vivo, each stage in morphogenesis emerges from a limited set of initial conditions. While cells self-organize efficiently at length scales around 100 µm, large changes in initial conditions can have a profound impact on growth outcomes^2,26–29^. To control initial conditions engineers have therefore developed platforms such as microwell arrays, microphysiological systems (or organs-on-a-chip), and 3D bioprinting that can be deployed together with biomaterials to further constrain self-organization^30–34^. Typically, however, the rheological properties required for fabricating these engineered systems are incompatible with those that are optimal for tissue morphogenesis, requiring that tissues are transferred between different materials (e.g. microwells) after fabrication or that their structure remains static within the device (e.g. microphysiological systems).

Among engineering platforms, embedded 3D bioprinting of saturated cell suspensions provides remarkable flexibility in generating initial conditions for tissue morphogenesis, including defining initial tissue size, composition, and geometry, both on-demand and in arbitrary microenvironments^35–44^. In principle, the capacity to “write” saturated cell suspensions directly into embedding materials holds the potential for controlling initial conditions and boundaries simultaneously. However, much of biomaterial design for 3D printing has focused on optimizing for the formation of complex and high-resolution geometries immediately following printing in a manner that sustains cell viability and simple behaviors like growth and motility. When the goal is to build tissues, however, active mechanical processes such as lumenization, inflation, compaction, cell sorting and folding that contribute to final tissue form are equally important^45–47^ (Fig. 1a). While rBMs support all of these behaviors, and Matrigel was previously employed as an embedded bioprinting material^45^, it has several critical limitations. Most notably, it behaves as a viscous fluid at 4 °C and quickly transitions to a soft hydrogel at 37 °C for cell culture. These regimes do not support embedded bioprinting except during a narrow period during gelation that is difficult to predict, limiting printing to ∼2–5 minutes and preventing generation of tissues of high complexity or in high-throughput.

**Figure 1.**
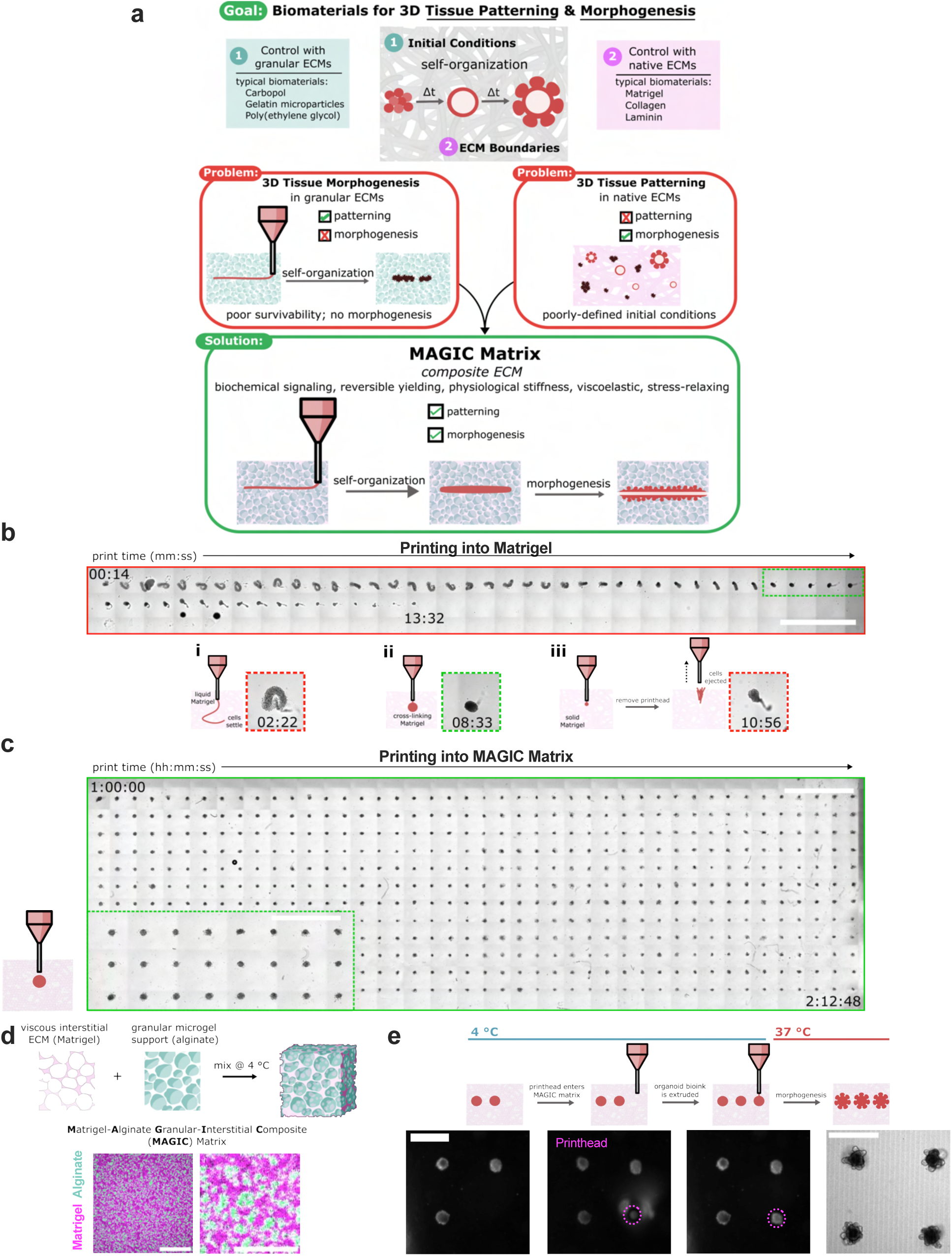
MAGIC extracellular matrices are embedded bioprinting materials that enable both patterning and morphogenesis of organoids. **a.** Biomaterials designed for tissue patterning, like some hydrogel microparticles, fail to support long-term tissue health or complex behaviors like morphogenesis. Biomaterials typically used for such behaviors include Matrigel and Collagen, but these materials do not mechanically support embedded bioprinting. MAGIC matrices exhibit a suite of biochemical and mechanical behaviors that, in sum, enable both complex patterning and morphogenesis of organoid bioinks. **b.** Organoid slurries printed into Matrigel while cross-linking at room temperature first distort and settle to the bottom of the dish (i) due to insufficient mechanical support, followed by a narrow regime (∼ 2 min) during which printing is viable (ii), and eventually are ejected or excluded as the hydrogel sets (iii). Insets show example of each regime. Scale bar = 4 mm. **c.** Organoid bioink printed into MAGIC matrix at 4 °C conforms to the desired morphology due its reversible yield-stress properties, enabling printing of many organoids (n = 528) over long times (t > 2h). MAGIC matrix was left in chilled conditions on the printer for 1 h before printing. Scale bar = 4 mm. Inset, zoomed view of organoids printed in MAGIC matrix. Scale bar = 2 mm. **d.** MAGIC matrix comprises of an inert alginate microgel granular support and a viscous basement membrane interstitium. Confocal microscopy images showing a standard MAGIC matrix composition using FITC-alginate microgels and NHS-labeled Matrigel. Scale bar = 100 µm. Zoomed view shows white pixels where fluorescent signals overlap, suggesting ECM in alginate microgels. Scale bar = 40 µm. **e.** Sequential brightfield images demonstrating MAGIC matrix bioprinting, in which the printhead enters cold MAGIC matrix, extrudes organoid slurry, and is then removed from the matrix, leaving a geometrically and spatially controlled feature behind. These organoid bioinks subsequently undergo self-organization at 37 °C to form mature organoids. Scale bars = 500 µm.

As an alternative, granular microgels are frequently applied as soft materials in embedded 3D bioprinting applications because they have been shown to support the structure of extruded materials after printing and are highly tunable in their properties^48,49^. A critical feature of these materials affecting print quality is reversible yield-stress behavior, in which the granular microgels yield in response to the printhead entering the bath and the extruded material, then recover to provide elastic support to the bioink once the nozzle is removed. However, most granular materials are not optimal for long-term cell health, and little is known about whether the properties necessary for bioprinting are compatible with complex tissue morphogenesis ^50,51^. Several groups have introduced interstitial matrices derived from natural ECM like collagen I with the goal of improving cell survival and dynamics^52,53^. However, the rheological properties and chemical composition of the resulting materials elicit cell behaviors that can be challenging to predict or counter to self-organization. For example, collagen I-containing ECM strain-stiffens and is not optimized to support epithelial growth and morphogenesis when compared to rBMs^45,54–57^. The ability of these materials to relax stress at long timescales and under physiological conditions is also poorly defined.

Here we quantify the relationship between granular biomaterial rheology and tissue self-organization to reveal that when controlling for storage and loss modulus at short timescales, the rate and extent of stress relaxation at long timescales determines the quantitative outcome of complex tissue morphogenesis. We apply this insight to design an optimized embedded bioprinting material comprising alginate microgels and interstitial Matrigel, termed <u>M</u>atrigel-<u>A</u>lginate <u>G</u>ranular-Interstitial <u>C</u>omposite (MAGIC) matrix. MAGIC matrices employ alginate microgels that are optically transparent and approximately cell-sized, which facilitated yielding behavior, high-fidelity embedded bioprinting, imaging, and organoid growth. Utilizing Matrigel as the interstitial material creates a switchable composite matrix that fluidizes upon shear at 4 °C to allow for long print times (≥2 h), but cross-links at 37 °C to create an environment that has similar rheology to pure Matrigel across several metrics. We find that MAGIC matrices provides remarkable tunability across properties including yield-stress, shear modulus, and stress relaxation at short and long timescales, including the ability to decouple loss tangent at short length- and timescales from stress relaxation over long timescales and large relative strains. To capitalize on the reproducibility, scalability, and automation afforded by 3D printing into MAGIC matrices, we designed a piezoelectric bioprinting platform that allows for full xyz-control, real-time imaging, and flexible scripted print geometries to control the initial conditions of organoid self-. The printhead enabled direct aspiration and extrusion of saturated cell slurries exceeding 10^8^ cells/mL, minimizing dead volume and required biomass, and supporting sub-100 µm print resolutions. Together, MAGIC matrix bioprinting led to nearly 100% organoid formation efficiency, dramatic improvements in inter-organoid homogeneity, orders-of-magnitude improvement in the statistical power of phenotypic assays, and enabled patterning of perfusable 3D microphysiological systems.

## Results

### MAGIC matrices can be tuned for optimal 3D patterning at 4 °C

Matrigel is the gold-standard biomaterial supporting complex morphogenesis of epithelial organoids. However, it does not support long bioprinting windows (Supplementary Movie 1). When extruding a cell-dense bioink as it warms from 4 °C to room temperature (as previously described^45^), Matrigel was initially too liquid to support printed shapes and filaments of cells settled to the bottom of the dish after extrusion (Fig. 1b, i). During a narrow span of ∼2 min during gelation, the bath supported printed shapes (Fig. 1b, ii), after which the gel became too elastic and cells were not extruded or ejected from the printing plane (Fig. 1b, iii). We therefore sought to design a reversible yield-stress granular material that supported embedded bioprinting at 4 °C but could be tuned to match the rheological properties of biomaterials such as Matrigel at 37 °C. For the granular phase we turned to alginate as an optically transparent biomaterial that is largely inert in mammalian culture. For these reasons, alginate has been employed in both bioprinting and organoid culture applications as a support medium and rheological modifier^19,37,58–60^. In addition, we employed microgels that were approximately cell-sized to facilitate self-organization, migration, and diffusion^52,61^ (Extended Data Fig. 1). We supplemented the alginate microgels with an interstitial matrix of Matrigel as its rich chemical composition including laminin and other basement membrane proteins, is necessary to support morphogenesis of most epithelial organoids. At 4°C, this material supported bioprinting of cell-dense bioinks for >2 h without compromising print fidelity (Fig. 1c). We termed this class of composite materials <u>M</u>atrigel-<u>A</u>lginate <u>G</u>ranular-Interstitial <u>C</u>omposite (MAGIC) matrix (Fig. 1d, e).

The composite biomaterial was tunable along several rheological parameters and in different temperature regimes to support bioprinting as well as proper tissue morphogenesis. To demonstrate this tunability we prepared several compositions spanning ranges likely to support both bioprinting and organoid culture and characterized their properties by shear rheology at 4 °C (relevant to bioprinting) and 37 °C (relevant to organoid culture). Preparations included undiluted jammed alginate microgels (AMGs); three different volume fractions of AMG slurry diluted in Matrigel at 2:1, 1:1, and 1:2 AMGs:Matrigel by added volume; and two different polymer weight fractions in the AMG preparation at 0.5 and 1 wt%. As previously reported, undiluted AMG slurries exhibited temperature-independent, but shear stress-dependent viscoelasticity, including yielding at ∼10 Pa shear stress for 0.5 wt% AMGs^58^ (Extended Data Fig. 2). In contrast, pure Matrigel exhibits significant temperature-dependence as it cross-links to form a hydrogel at physiological temperature (Extended Data Fig. 2).

All compositions of MAGIC matrix demonstrated reversible yield-stress behavior at 4 °C, suggesting their utility as embedded bioprinting materials (Fig. 2a). At rest, they behaved as viscoelastic solids, with G’ greater than G’’, in contrast to pure Matrigel (Extended Data Fig. 2). However, as shear stress was increased, G’’ overtook G’, indicating microgel yielding and rearrangement. These materials remained as reversible yield-stress fluids for as long as they were kept cold, suggesting their ability to support long print times. The specific yield-stress of each composition was measured by unidirectional shear tests and fit well to a Herschel-Bulkley exponential model (Extended Data Fig. 2). Despite the similarity of the interstitial ECM, varying MAGIC matrix composition tuned yield-stress values over an order of magnitude (Fig. 2b). Viscoelasticity and yield-stress were increased when using a viscous interstitial material such as Matrigel as opposed to diluting in equivalent volumes of cell culture media (Extended Data Fig. 2), highlighting the importance of the interstitial phase in granular materials at 4 °C^62^. Together, these results imply that the material can be engineered to fit the needs of a particular printing regime.

**Figure 2.**
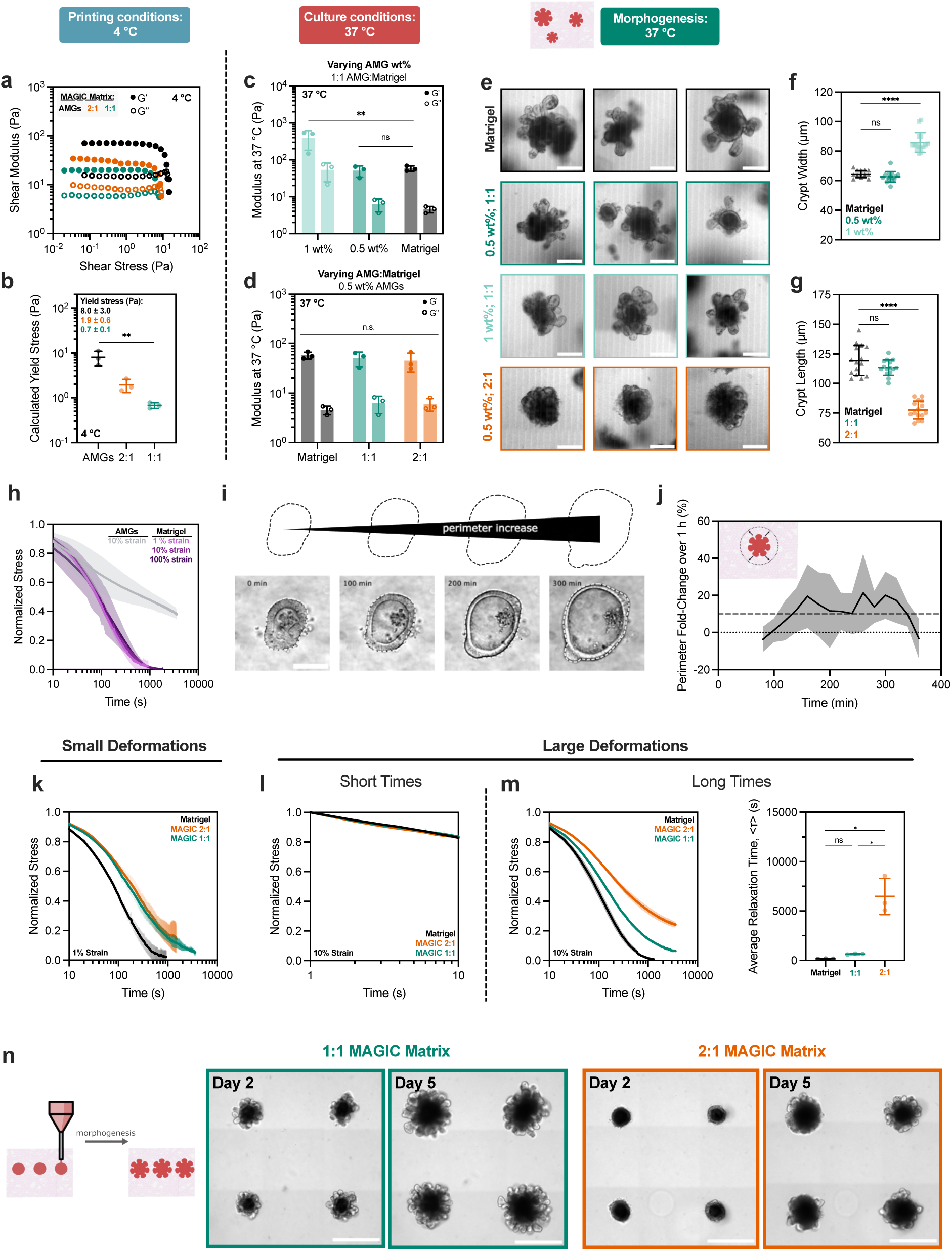
Rheological properties of MAGIC matrices including low stiffness and stress relaxation drive gold-standard morphogenesis. **a.** At 4 °C, oscillatory amplitude sweeps at 1 Hz reveal that various MAGIC matrix compositions behave as yield-stress materials, indicated by G’ and G’’ cross-over. **b.** MAGIC matrices behave as Herschel-Bulkley fluids at 4 °C with yield-stresses calculated using a power law model. **c.** Storage and loss moduli at 1 Hz and 1% strain of MAGIC matrices at 37 °C prepared from 1 wt% or 0.5 wt% AMGs. **d.** Storage and loss moduli at 1 Hz and 1% strain of MAGIC matrices at 37 °C prepared using different volume ratios of 0.5 wt% AMGs:Matrigel. **e.** Mouse intestinal organoids grown in pure Matrigel (top row) and MAGIC matrices of several compositions (lower rows) five days after seeding. Scale bars = 200 µm. **f.** Quantification of organoid crypt width as a function of matrix composition. **g.** Quantification of crypt length as a function of matrix composition. **h.** Normalized stress relaxation curves for undiluted 0.5 wt% AMG slurry at 10% strain or Matrigel over 1 h at 1%, 10%, or 100% strain. **i.** Brightfield images and segmented cartoons simulating organoid growth over time in Matrigel. Scale bar = 100 µm. **j.** Measurement of material strain at the organoid-ECM interface over time quantified using segmented organoid perimeter. The dashed line at 10% strain represents the strain value used for most stress relaxation measurements. **k.** Normalized stress relaxation curves for Matrigel or MAGIC matrices over 1 h at 1% strain. **l.** Normalized stress relaxation curves for Matrigel or MAGIC matrices over 10 s at 10% strain. **m.** Normalized stress relaxation curves for Matrigel or MAGIC matrices over 1 h at 10% strain (left) and quantification of average relaxation time for each matrix using a stretched exponential model (right). **n.** Organoid array bioprinting experiments demonstrating similar impacts on crypt morphogenesis when MAGIC matrix compositions are employed as bioprinting support baths. Scale bars = 500 µm. For all rheological experiments, data shown are mean ± SD from n = 3 independently prepared replicates. For crypt length measurements, data shown are mean ± SD on the median n = 15 crypts from ≥10 organoids per matrix condition. For ECM strain during organoid growth, data shown are mean ± SD of n = 9 organoids. * = p ≤ 0.05; ** = p ≤ 0.01; **** = p ≤ 0.0001; ns = not significant determined by one-way ANOVA with Tukey’s multiple comparisons (**b**, **c, m**) or Dunnett’s multiple comparisons (**f, g**).

### Independent control over modulus and stress relaxation in MAGIC matrices uncovers the contributions of ECM rheology to crypt morphogenesis

To assess MAGIC matrix behavior under physiological conditions, we next measured viscoelastic properties following cross-linking at 37 °C. At 0.5 wt% alginate, the storage and loss modulus of the composite matrix were similar to that of pure Matrigel (Fig. 2d). This was true across a range of MAGIC matrix compositions, implying that for these formulations, the rheological properties of the composite matrix may be dominated by cross-linked Matrigel. In contrast, substituting 0.5 for 1 wt% AMGs significantly increased storage and loss moduli (Fig. 2c). These analyses suggested that the various preparations of 0.5 wt% MAGIC matrices had rheological properties and interstitial medium that were similar to Matrigel, and therefore, may equivalently support tissue growth and morphogenesis.

We evaluated the performance of these ECM compositions in morphogenesis assays using mouse duodenal organoids. Intestinal organoid crypt budding and lumen expansion is highly sensitive to matrix mechanics^11,15,63,64^. We therefore measured crypt width and length across all matrix formulations and in comparison to Matrigel in manual dome culture. Consistent with their increased stiffness, all organoids in MAGIC matrices containing 1 wt% AMGs formed crypts with altered structures, with shorter and wider morphologies than Matrigel controls (Fig. 2e, Extended Data Fig. 3). After five days in culture, all 1 wt% matrices yielded organoids with shorter and wider crypts (Fig. 2f). In contrast, MAGIC matrix compositions employing 0.5 wt% at 50% volume fraction or lower AMGs were indistinguishable from those grown in pure Matrigel (Fig. 2e, f, g). As with pure Matrigel, the 0.5 wt% MAGIC matrix formulations were notably softer than many traditional bioprinting and biomaterial scaffolds^17^. Moreover, composite matrices formulated with both with 25% Matrigel and 1 mg/mL collagen I were significantly stiffer than 0.5 wt% MAGIC matrices and gave rise to organoids with minimal or no crypts^58^ (Extended Data Fig. 4). These results highlight the importance of soft, non-fibrillar basement membrane proteins for supporting gold-standard epithelial morphogenesis.

We were surprised to find that formulations of MAGIC sharing 0.5 wt% microgels but having different volume fractions of Matrigel had significantly different crypt morphologies despite their nearly identical storage and loss moduli. Specifically, 2:1 AMGs:Matrigel had significantly shorter and wider crypts compared to 1:1 MAGIC and Matrigel (Fig. 2g, Extended Data Fig. 3). These findings were not a consequence of Matrigel dilution, indicating that potential differences in the concentration of the interstitial ECM did not contribute to these observations (Extended Data Fig. 3).

To begin to understand which properties of these two material formulations that were contributing to crypt phenotype, we noted that intestinal organoids undergo large morphological changes over hour timescales during morphogenesis, including crypt budding and elongation as well as lumen inflation and collapse cycles^64,65^, all of which induce large mechanical strains on the surrounding hydrogel. The magnitude and rate of these volumetric changes were previously shown to be associated with major differences in morphogenesis^15,63^. Additionally, stress relaxation over these timescales affects single-cell behaviors^18,19,66–68^ and is required for symmetry breaking events at the tissue-scale^17,60,69–72^. Thus, we hypothesized that crypt morphology may be quantitively linked to stress relaxation rate and magnitude at long timescales and large strains, even when the response of the materials at short length- and timescales was identical. This hypothesis was not previously testable, as prior investigations of stress relaxation used materials whose stress relaxation rate was correlated with the loss tangent at short timescales^18,60,66–70^.

To test this hypothesis, we first measured stress relaxation of Matrigel over 1 h and at variable strain. Notably, Matrigel completely relaxed accumulated stresses in ∼15 min for a variety of strain magnitudes (Fig. 2h). In contrast, undiluted AMG slurry did not fully relax accumulated stress. We next investigated the relevant length scale of stress relaxation for intestinal organoid morphogenesis by quantifying organoid perimeter growth over time^73^. Over 1 h segments we observed an average perimeter expansion of ∼10% (Fig. 2i, j). Importantly, these changes in organoid size occur over timescales much longer than standard biomaterial measurement regimes (∼1 Hz). We then compared stress relaxation properties of 1:1 and 2:1 MAGIC matrix formulations given that these compositions had matched storage and loss moduli but exhibited striking differences in crypt morphology. For small deformations (approximated as 1% strain), the 1:1 and 2:1 MAGIC matrices dissipated stress nearly identically over both short and long timescales (Fig. 2k). While both formulations dissipated stress more slowly than pure Matrigel, they ultimately dissipated 80-90% of accumulated stress after 1 h. For larger deformations relevant to organoid morphogenesis (10% strain), stress relaxation of both MAGIC formulations and Matrigel occurred nearly identically at short timescales (≤ 10 s) (Fig. 2l). However, over long times (10% strain over 1 h), 2:1 MAGIC matrix relaxed stress significantly more slowly than both Matrigel and 1:1 MAGIC, providing a potential explanation for their differential effects on organoid morphogenesis (Fig. 2m). Critically, 1:1 MAGIC matrices dissipated ∼95% of internal stresses after 1 h, with as little as ∼0.2 Pa of residual stress remaining in the material.

To quantify the relevant timescale of stress relaxation we fit stress relaxation profiles using a stretched exponential function^68^ (Supplementary Information, Extended Data Fig. 4). At 10% strain, Matrigel and 1:1 MAGIC matrices had the shortest average relaxation time, followed by an increased relaxation time 2:1 MAGIC matrices (Fig. 2m). This bulk property of the materials was further explored at the micron scale by performing nanoindentation to measure stress relaxation, where we observed impeded relaxation in 2:1 MAGIC matrix compared to 1:1 MAGIC and Matrigel (Extended Data Fig. 4). Notably at this scale, 1:1 MAGIC matrix and Matrigel relaxed stress at a similar rate and magnitude. A previous study found that fully synthetic matrices with a small degree of stress relaxation could promote symmetry breaking toward crypt morphogenesis, although these organoids formed shorter and smaller crypts than those in Matrigel^21^. In contrast, MAGIC matrices of all formulations dissipated ≥70% of stress over 1 h, with formulations having the fastest stress relaxation promoting crypt morphogenesis indistinguishable from Matrigel. These findings were further supported by creep-recovery measurements at 37 °C using 10 Pa applied shear stress over 10 minutes to simulate the forces and timescales of processes such as lumen expansion and crypt budding^63,74^ (Extended Data Fig. 5). The creep response was a strong function of material composition, with Matrigel and MAGIC matrices comprising soft (0.5 wt%) AMGs exhibiting the greatest strain rate and highest plasticity.

To investigate whether decoupling of short and long length- and timescale stress relaxation was a general feature of MAGIC matrices, we repeated these measurements in formulations spanning three alginate wt% in the microgels and 1:1 and 2:1 volume fractions. Indeed, for all compositions, stress relaxation rate and magnitude could be tuned for materials with matched storage and loss moduli (Extended Data Fig. 6). While storage and loss moduli were a function of wt% in the AMG fraction, they were independent of volume fraction for 1:1 and 2:1 MAGIC matrix formulations. Furthermore, the volume fraction of AMG dictated the stress relaxation profiles independent of wt% in the microgels, with 2:1 matrices relaxing at a slower rate than 1:1. Interestingly, stiffer microgels corresponded to decreased average relaxation time (Extended Data Fig. 6). While the loss tangent and relaxation properties are often coupled for homogenous or non-granular viscoelastic materials^19,56,66–70,75^ (Extended Data Fig. 6), these data suggest that these may be de-coupled at larger deformations for granular materials with viscous interstitium, possibly due to microgel yielding and rearrangement. These results suggest that tuning microgel packing and geometry may present unique opportunities to independently investigate the roles of loss tangent and stress relaxation for small and large deformations on 3D tissue behaviors.

To establish that these properties of MAGIC matrix translated to a 3D bioprinting context where cells are directly extruded into the support bath, we measured the impact of stress relaxation on crypt morphogenesis using a bioink derived from a saturated suspension of dissociated intestinal organoids (see next section). Crypts appeared more slowly in 2:1 MAGIC matrix compared to 1:1, suggesting that rate of crypt morphogenesis may also be impacted by stress relaxation (Fig. 2n). Taken together, these results demonstrate that for materials that are matched in viscoelasticity at the short length- and timescales typical of rheological measurements, differences in mechanical properties at long timescales and large strains can profoundly impact complex biological processes like morphogenesis.

### A piezoelectric printhead enables precise, automated direct “writing” of dense cell slurry bioinks

Organoid morphogenesis can be highly uniform when using appropriately dynamic boundaries and carefully controlling for initial conditions, for example by growing organoids from cell aggregates of similar initial size, geometry, and composition^45,52,76,77^. However, current platforms for controlling initial conditions rely on materials that do not support morphogenesis, requiring transfer of organoids into appropriate matrices, or limiting flexibility in the geometry and configuration of initial conditions. We therefore focused on developing bioprinting modalities for controlling initial conditions that use MAGIC matrices together with bioinks prepared as single-cell slurries of dissociated organoids at tissue-like densities ≥10^8^ cells/mL^78–80^. We designed a printhead that could directly aspirate small volumes with high precision, fast pressure ramps, and without excess loss due to tubing and fluidics (i.e. “dead volume”). We used a piezoelectric actuator that is mechanically coupled to a fluid-filled cavity via a polyether ether ketone (PEEK) diaphragm (Fig. 3a)^81^. Voltage applied to the piezoelectric material to be translated into precise expansion or contraction of the diaphragm, and therefore, aspiration and dispensing of precise volume of cell suspension. The printhead was controlled with motorized arms and mounted on a standard commercial microscope to facilitate alignment and live analysis of prints. Using this piezoelectric extrusion bioprinting setup, it was possible to print large tissue arrays comprising >100 individual organoids, beginning with fewer than 10^6^ cells as a saturated slurry.

**Figure 3.**
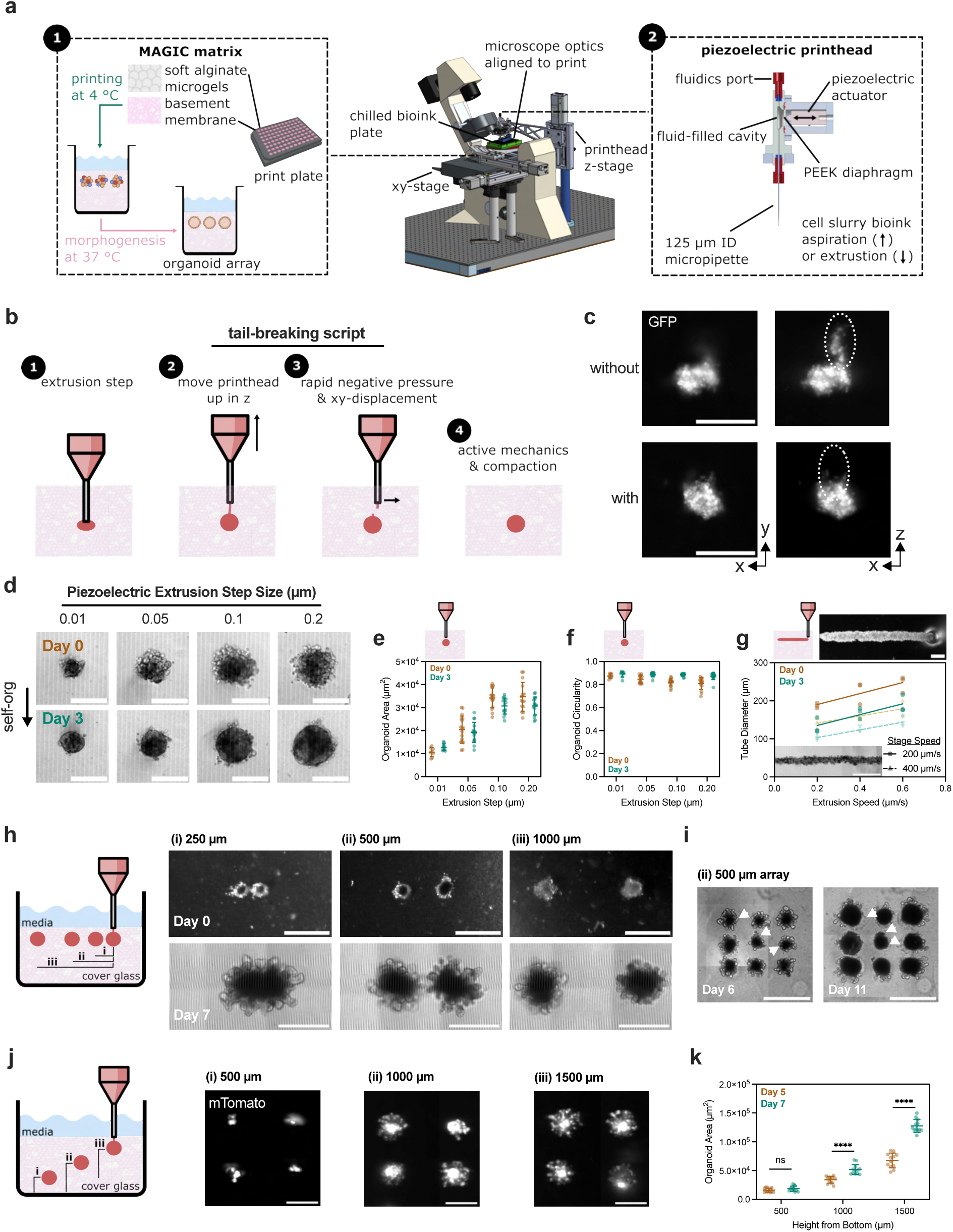
MAGIC matrix bioprinting utilizes a piezoelectric printhead to precisely aspirate and extrude cell slurry bioinks and can be programmed to generate organoid arrays. **a.** Schematic illustrating the MAGIC matrix bioprinting platform, including the benefits of MAGIC matrix (left) and the piezoelectric printhead (right). **b.** The piezoelectric printhead and precise xyz-control tolerate rapid pressure ramps and print plate movement, enabling scripts such as tail-breaking to improve print fidelity of viscous cell slurry bioinks. **c.** Representative fluorescent images of bioprinted spheroids with and without a tail-breaking script enabled by piezoelectric bioprinting through rapid changes in applied voltage and printhead position. Scale bars = 200 µm. **d.** Representative brightfield images of bioprinted Caco-2 tissues at day 0 and day 3 post-print as a function of extrusion step size controlled via applied voltage. Data are representative of at least n = 9 individual tissues per extrusion step condition, scale bars = 200 µm. **e.** Organoid area and **f.** circularity measured using max intensity projections of confocal z-stack images of GFP-expressing Caco-2 cell slurries. At both day 0 and day 3, organoid area significantly depends on extrusion step size until beyond 1.0 µm as determined by one-way ANOVA with Tukey’s multiple comparisons. Data shown are mean ± SEM of n ≥ 9 individual tissues per extrusion condition. At day 3, organoid circularity does not depend significantly on extrusion step size as determined by one-way ANOVA. **g.** Quantification of bioprinted tube diameter at day 0 and day 3 post-printing as a function of both stage translation speed and extrusion step speed. Fit demonstrates that initial and final tube diameter are approximately linear functions of extrusion step speed for a given stage translation speed. Above, representative brightfield image of organoid tube during print; scale bar = 200 µm. Inset shows representative tube exhibiting signs of lumenization 3 days after printing; scale bar = 500 µm. Data shown are n = 3 bioprinted tubes per condition. **h.** Bioprinted mouse intestinal organoid pairs printed with 250, 500, or 1000 µm center-to-center organoid spacing. Tissues printed close together (∼75 µm edge-to-edge) fuse (i), whereas tissues printed far enough apart (>∼300 µm edge-to-edge) don’t fuse (ii, iii). Scale bars = 500 µm. **i.** Organoids printed in arrays with 500 µm pitch lack crypts between day 6 and 11. Arrowheads indicate crypts that formed close to the neighboring organoids that are gone by day 11. Scale bars = 500 µm. **j.** Maximum intensity projections of intestinal organoid arrays bioprinted at different depths within MAGIC matrix (500 µm, 1000 µm, or 1500 µm from the cover glass). Scale bars = 500 µm. **k.** Organoids printed deeper in the matrix (i) do not significantly grow between days 5 and 7 post-print compared to organoids printed closer to the media interface (ii, iii). Data shown are mean ± SD of n = 12 organoids per condition; ns = not significant, **** = p < 0.0001 determined by non-parametric t-test between day 5 and day 7.

High-viscosity ink droplets such as cell suspensions are challenging to print in yield-stress fluids due to inertial, cohesive, and shear forces between the ink and matrix^82,83^. For example, cohesive interactions between cells in the slurry causes the ink to be pulled as a continuous tail from the tip as it is removed from the bath, generating a structure reminiscent of capillary bridging^84,85^. To combat this, we leveraged a rapid voltage switch on the piezo printhead to “break” this tail by applying a small negative pressure and xz-displacement (Fig. 3b, c). We then applied this aspiration, extrusion, and tail-breaking program to script an automated spheroid bioprinting array. We first demonstrated that a Caco-2 cell slurry bioink could be delivered through a 125 µm ID nozzle to create spheroids of customizable dimensions (Fig. 3d). Spheroid area was a linear function of extrusion step (Fig. 3e).

After bioprinting and self-organization in MAGIC matrices, Caco-2 spheroids underwent efficient lumenization^86^ (Fig. 3d). While the initial geometry of extruded cell volume sometimes deviated from spherical, the high active surface tensions of these living inks rapidly correct surface irregularities through self-organization (Fig. 3f). Using scripting, cell slurries could also be printed as cylinders. Cylinder diameter could be tuned using both extrusion rate and stage speed, providing multiple engineering controls (Fig. 3g). Tube diameter was an approximately linear function of extrusion rate for the two stage speeds tested. Bioprinted Caco-2 cylinders self-organized similar to spheres, undergoing compaction followed by lumenization while maintaining the high aspect ratio programmed by their unique initial conditions (Fig. 3g, inset). Therefore, the piezoelectric printhead affords high volume precision, working with small volumes of precious cellular inks, and flexible control over bioink behavior by rapidly changing applied voltage. To our knowledge, this is the first demonstration of applying a piezoelectric actuator to control direct aspiration and extrusion of dense cell slurries.

We explored the generality of bioprinting into MAGIC matrices by preparing mouse small intestine organoid arrays, which undergo compaction, lumenization, symmetry breaking, and subsequent morphogenesis to form crypts over several days in culture (Supplementary Movie 2). A significant contributor to tissue and organoid heterogeneity in traditional ECM dome cultures is the disparity in microenvironmental conditions such as media access or inter-organoid spacing^87,88^. To assess the impact of initial spatial conditions on organoid growth, we systematically varied either inter-organoid spacing or z-depth within the MAGIC matrix support bath using organoids printed with an initial average diameter of 200 µm. Organoid pairs printed with 250 µm centroid-to-centroid spacing came into direct contact as they grew and fused over time, leading to one large organoid (Fig. 3h), indicating that MAGIC matrix does not interfere with matrix rearrangements necessary for tissue-tissue fusion^64,76^. Organoids spaced 500 or 1000 µm apart remained distinct and appeared to undergo normal morphogenesis through at least 7 days of culture. To examine the effect of inter-organoid spacing on growth and morphogenesis we printed 3 x 3 arrays with 500 µm spacing. Notably, after 11 days in culture, crypts only appeared at the periphery of the organoid array, while the central organoid remained compacted and lost crypts (Fig. 3i). We hypothesize that this could be caused either by nutrient depletion by the outermost organoids or autocrine gradients^89–92^. Printing depth also had a profound effect on organoid phenotype, with organoids printed closer to the ECM-media interface growing the largest and forming the most crypts (Fig. 3j). Organoids printed too far from the ECM-media interface did not significantly grow over the same time (Fig. 3k). Together, these results highlight the importance of initial conditions and local microenvironment on organoid morphogenesis and imply that these effects could impact experimental results when working with organoids in manual 3D culture.

### MAGIC matrix enables the self-organization of many organoid types following bioprinting

Having established initial conditions supporting controlled intestinal organoid growth, we next assessed whether organoid arrays from these and other developmental lineages exhibited expected mechanisms of self-organization when printed and cultured in MAGIC matrices, including processes like lumen formation, folding, budding, sprouting, and sorting. For mouse duodenal organoids, live imaging of a membrane-localized tdTomato reporter revealed crypt budding and protrusion within 2–3 days after printing in 96-well microplates (Extended Data Fig. 7). In these folded epithelial regions, *Lgr5*+ stem cells were appropriately positioned toward the base of the structures (Fig. 4a). Fixed intestinal organoids stained positive for mature epithelial cell types, including Paneth cells (lysozyme) and enteroendocrine cells (chromogranin-A). Cohesive and continuous cell borders were marked with E-cadherin. 3D reconstruction of organoids in MAGIC matrix revealed that crypts radiated in all directions from the organoid lumen. Like Caco-2 cells, cell slurries derived from mouse duodenal organoids also self-organized efficiently when printed in a cylindrical format^45,76,93^. Printed cords compacted, lumenized and began folding into crypts which formed radially around the long axis of the intestinal tube within 2–3 days after printing. Cell patterning in tubular crypts were similar to those in spherical intestinal organoid arrays (Fig. 4a).

**Figure 4.**
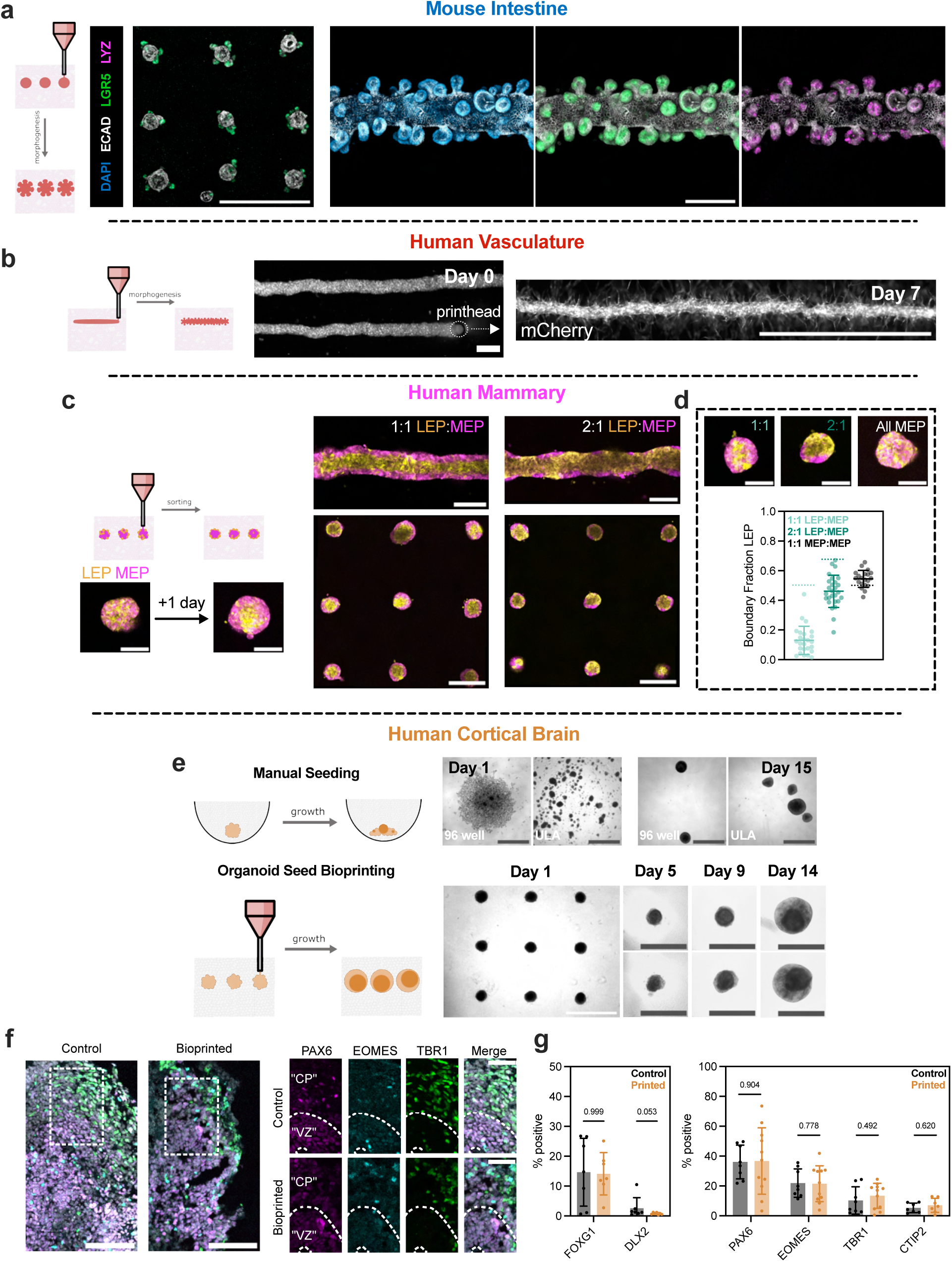
MAGIC matrices promote self-organization of bioprinted tissues derived from the three primary germ layers. **a.** Left, maximum intensity projections of a representative *Lgr5*-GFP and ECAD-stained intestinal organoid array 3 days after bioprinting. Scale bar = 1 mm. Right, maximum intensity projections of DAPI- (left), *Lgr5*-GFP (middle), and LYZ (right) stained intestinal organoid tubes 3 days after bioprinting. Scale bar = 200 µm. **b.** Representative live images of bioprinted mCherry-expressing HUVEC tubes during printing (day 0) and following self-organization (day 7). Tubes over 2 mm long could be printed, with signs of vascular sprouting. Scale bars = 200 µm (day 0) and 1 mm (day 7). **c.** Maximum intensity projections of bioprinted spheroid arrays and tubes of human mammary epithelial cell (HMEC) organoids of different luminal and myoepithelial compositions. Organoids were allowed to sort for one day following printing. Scale bars = 500 µm for arrays and tubes; scale bars = 200 µm for individual spheroids. **d.** Representative live images and quantification of luminal cell boundary occupation in bioprinted organoids as a function of composition. Dashed lines represent expected boundary occupancy for mechanically equivalent cells^77^. Data shown are mean ± SD for n ≥ 20 organoids analyzed per composition. Scale bars = 200 µm. **e.** Comparison of manually seeded and bioprinted induced pluripotent stem cell-derived human cortical brain organoids. Brightfield images of manually seeded cortical brain organoids in 96 or ultra-low attachment (ULA) well plates (top) or bioprinted arrays (bottom) over time. Scale bars = 1 mm (array) or 200 µm (manually seeded or individual bioprinted organoids). **f.** Left, 18 µm maximum intensity projections of control or bioprinted cortical organoid sections stained for cortical identity and neuronal differentiation. Scale bars = 100 µm. Right, staining for cortical cell types. “CP” = cortical plate; “VZ” = ventricular zone. Scale bars = 50 µm. **g.** Quantification of cortical identity (left) and neuronal differentiation (right) between bioprinted and manually seeded cortical brain organoids. Data shown are mean ± SEM of 1 quantified cryosection of n ≥ 3 organoids per marker from 2 separate differentiations; numbers shown are p values of multiple Mann-Whitney tests.

Organoids of multiple developmental lineage that form through different programs of self-organization behaved as expected following MAGIC matrix bioprinting. For example, we bioprinted organoid arrays from mouse submandibular salivary gland, an ectodermal tissue which undergoes budding morphogenesis^94^ (Extended Data Fig. 7). These tissues exhibited characteristic multi-lobular structures within 3 days of printing and expressed ductal and basal epithelial markers, keratins 8 and 14, respectively. We also printed vascular cords from cell slurries derived from human umbilical vein endothelial cell (HUVEC) cultures. After 7 days in culture, these vascular cords sprouted microvessels when printed into MAGIC matrix compositions supplemented with 1 mg/mL collagen I^93^ (Fig. 4b, Supplementary Movie 3). Bioprinted vessels could be tuned in length and width over a wide range up to and beyond the centimeter-scale (Supplementary Movie 4). Using mouse embryonic fibroblasts, we demonstrated that matrix-in-matrix patterning between organoids could also control their collective migration along pre-defined tracks using a 200 µm ID nozzle to print ECM (Extended Data Fig. 7). In addition to lumen formation and tissue folding, cell sorting is a common mechanism of self-organization that derives from differences in cell-cell and cell-ECM interfacial interactions among different cell lineages^6,26^. We evaluated the efficiency of cell sorting by printing spheroid arrays of patient-derived human mammary epithelial cells (HMEC) organoids which comprise both the luminal cell (LEP) and myoepithelial cell (MEP) lineages^77,95^ (Fig. 4c). Printed tissues robustly sorted across ∼100 µm distances to form bilamellar structures within one day, with the continuity of MEP coverage in the outer layer proportional to MEP composition in the organoid (Fig. 4d). HMECs printed as tubes also sorted robustly and maintained their initial tissue tubular geometry. In contrast, MEP-only spheroids and tubes showed no evidence of sorting. Thus, MAGIC matrix bioprinting is compatible with a variety of self-organization mechanisms, including sorting, lumenization, migration, and tissue folding.

The most common approach for defining the initial conditions of organoid culture is to aggregate dissociated tissue or stem cells using hanging droplets or low attachment plates^1^. However, spheroid formation using these methods can occur along complex or inefficient trajectories and are strongly dependent on the ability of the cells to aggregate and compact. Consequently, the process of aggregation often includes a field of unincorporated or dead cells at the periphery of the organoids. It is unclear how these complex aggregation dynamics might impact downstream morphogenesis (Fig. 4e). We reasoned that MAGIC matrix bioprinting could enhance aggregate formation by forcing cells to interact in a common volume. To test this idea, we explored the feasibility of bioprinting human induced pluripotent stem cell (iPSC)-derived forebrain organoids^96,97^. We dissociated brain organoids after 7 weeks of differentiation and culture to form a cell slurry bioink that was printed into both MAGIC matrix or a pure AMG bath and assayed for self-organization of cellular rosettes and allocation of expected cell types across neural identities. Cortical brain organoids printed into MAGIC matrix exhibited the expected sprouting behavior that occurs in rBM gels, as well as neuroepithelial bud formation, but lacked neuroectoderm (Extended Data Fig. 8)^98,99^. In contrast, cortical brain organoids printed in pure AMG slurry formed dense spheroids without protrusions and exhibited spatially organized neuroectoderm (Fig. 5e). These organoids were positive for the dorsal forebrain marker *FOXG1* and negative for the ventral forebrain marker *DLX2*, with ∼15% dorsal identity <1% ventral identity, comparable to iPSC-derived cortical organoids aggregated with low attachment wells and not bioprinted (Fig. 5f, g). Furthermore, the bioprinted cortical organoids demonstrated characteristic self-organization with neural progenitors surrounding ventricular zone-like regions (PAX6+, ∼35%), intermediate progenitor cells (EOMES+, ∼20%) and deep layer excitatory neurons (TBR1+, ∼10%; CTIP2+, ∼5%) extending radially out from the apical surface (Fig. 5f). Overall, bioprinted organoids showed similar cell proportions to traditional cortical organoids. Together, these results demonstrate the efficiency of aggregate formation, validate organoid bioprinting for iPSC-derived tissues, and lay the groundwork for generating more complex organoid interfaces such as assembloids^100^.

**Figure 5.**
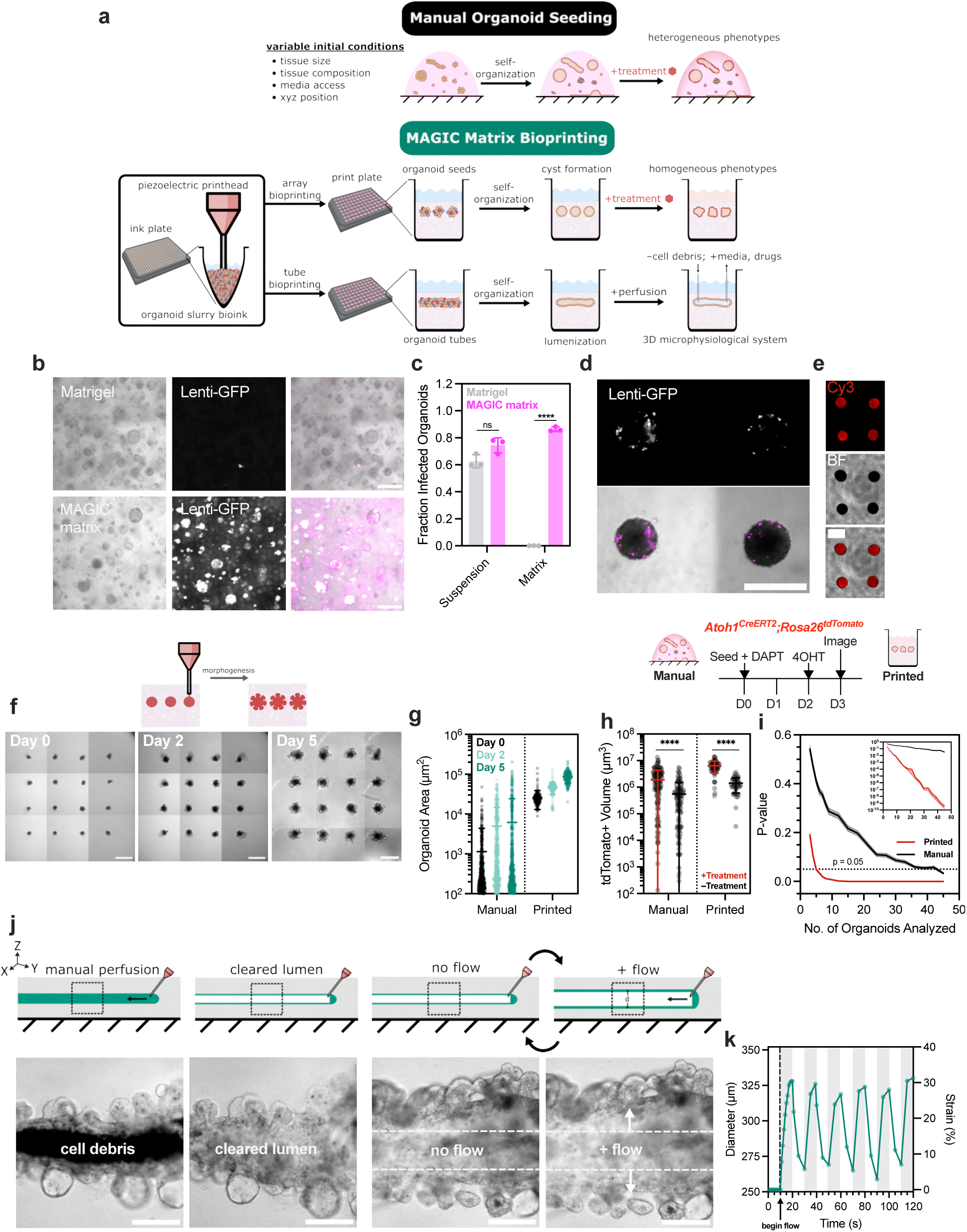
MAGIC matrix bioprinting enables generation of high-throughput organoid arrays with improved statistical power and perfusable 3D microphysiological systems. **a.** Traditional manual methods of seeding organoids lead to heterogeneity in organoid growth and morphogenesis due to heterogeneity in starting tissue size, composition, and microenvironment. By controlling for initial conditions such as cell number, media access, and organoid spacing, bioprinting platforms facilitate rapid generation of reproducible organoid arrays or freeform 3D microphysiological systems**. b.** Live images of triple-negative breast cancer (TNBC) patient-derived organoids transduced overnight with GFP-expressing lentivirus while seeded in Matrigel (top) or MAGIC matrix (bottom). Scale bars = 200 µm. **c.** Fraction of GFP+ organoids transduced in suspension before seeding or transduced after seeding for either ECM composition. Data shown are mean ± SD of n = 3 replicate ECM conditions; ns = not significant, **** = p < 0.001 determined by non-parametric t-test between ECM conditions. **d, e.** Live images of bioprinted TNBC organoids transduced with GFP-expressing lentivirus overnight after printing (**d**) or transfected using Lipofectamine and Cy3-conjugated single-stranded non-coding small RNA for 24 h, 3 days after printing (**e**). Scale bars = 500 µm. **f.** Live imaging of bioprinted intestinal organoid arrays following printing and after 2 and 5 days in culture. Scale bars = 500 µm. **g.** Organoid area over time measured by segmentation for manual or bioprinted intestinal organoids; data shown are mean ± SD of n ≥ 190 organoids per time point. **h.** Top, experimental outline of phenotypic assay for inhibition of gamma-secretase. Red fluorescence indicates *Atoh1+* secretory progenitors. Bottom, total red fluorescence volume per organoid in treated and untreated conditions from live images of bioprinted and manually seeded organoids treated with and without gamma-secretase inhibitor. For bioprinted arrays, data shown are mean ± SD of n = 45 organoids per condition. For manually seeded organoids, data shown are mean ± SD of n ≥ 135 organoids per condition. **** = p < 0.0001 determined by non-parametric t-test. **i.** Bootstrapping analysis of statistical significance between treated and untreated conditions for either bioprinted or manually seeded organoids as a function of number of paired comparisons. Inset shows statistical significance approaches zero (< 10^-9^) for bioprinted organoids using an equivalent number of comparisons as it takes manually seeded organoids to approach p = 0.05. **j.** Brightfield images of bioprinted intestinal organoid tubes that are manually perfused with a glass capillary attached to a micromanipulator and flushed to remove cell debris and access the lumen. Scale bars = 200 µm. **k.** Quantification of tube diameter and resulting strain upon application and removal of fluid flow. Gray bars correspond to times when fluid flow was applied.

### Generation of high-throughput bioprinted organoid arrays for assay development

Similar to microwell methods, bioprinted organoid arrays hold great promise to increase the reproducibility and sensitivity of high-throughput assay development using CRISPR and drug screens^31^. However, bioprinted arrays have the added benefit of precise control over tissue geometry and other initial conditions (Fig. 5a). We compared organoid arrays prepared using MAGIC matrix bioprinting to traditional microwell aggregation. Arrays prepared using MAGIC matrix bioprinting had decreased variance in initial tissue size and improved spheroid circularity compared to microwells. Using the 125 µm nozzle and standard bioink densities, they were less effective in seeding very small organoids comprised of ∼100 cells each (Extended Data Fig. 9), presenting an opportunity to optimize bioprinting of small organoids by tuning these parameters. To explore the utility of MAGIC matrix for assays requiring genetic perturbations, like CRISPR screens, we transduced triple-negative breast cancer (TNBC) patient-derived organoids using lentivirus encoding expression of H2B-GFP (Fig. 5b). TNBC organoids showed no difference in infection efficiency when transduced in suspension and then plated in Matrigel or MAGIC matrix domes (Fig. 5c). However, when transducing TNBC organoids through the ECM domes by including lentivirus in the media after plating and gel cross-linking, infection efficiency was dramatically increased in MAGIC matrix, with nearly 90% GFP+ organoids compared to minimal-to-no infection in Matrigel. We hypothesize this is due to improved diffusivity or low protein binding afforded by low wt% (and therefore large pore size) AMGs in the composite support. Cells within TNBC organoids were also successfully transduced by including lentivirus in the media after bioprinting (Fig. 5d). TNBC organoid arrays were also highly amenable to transfection using Lipofectamine, showing strong RNA uptake as a function of both transfection time and amount of RNA delivered (Fig. 5e, Extended Data Fig. 9).

Achieving regularity among organoids with complex morphologies is a key challenge in developing organoid-based assays, as it impacts the response of the organoids to genetic, mechanical and chemical perturbations^31,101,102^. Structural and phenotypic heterogeneity among manually seeded organoids include differences in organoid size, morphological features such as number of crypts, and position across multiple z-planes, all of which can obscure subtle phenotypes or lead to altered phenotypes that are only revealed after analyzing dozens to hundreds of individual organoids, typically using imaging-based screens. For example, crypt morphogenesis in mouse small intestinal organoids is highly sensitive to the chemical and mechanical microenvironment^11,73^. We therefore quantified the timing and regularity of crypt morphogenesis after bioprinting in MAGIC matrix compared to manual culture methods. Over the course of 5 days, bioprinted organoids synchronously self-organized, initially into lumenized cysts that further underwent budding morphogenesis to form crypts (Fig. 5f). By contrast, manually seeded organoids showed far more heterogeneous sizes and morphologies over the same timeframe (Fig. 5g). Furthermore, for a given time post-seeding, bioprinted organoid arrays underwent more extensive and uniform morphogenesis, exhibiting a greater number of crypts with decreased variance compared to manually seeded organoids^103^. Organoids grown in manually seeded cultures using MAGIC matrix as opposed to pure Matrigel showed no difference in crypt number, indicating more uniform initial conditions specifically contribute to improved maturity (Extended Data Fig. 9).

### Bioprinted organoid arrays dramatically improve assay statistical power

The uniformity and reproducibility afforded by MAGIC matrix bioprinting could dramatically improve the sensitivity of phenotypic assays while reducing required input material. We test this idea by treating intestinal organoids with a gamma-secretase inhibitor, N-[N-(3,5-Difluorophenacetyl)-L-alanyl]-S-phenylglycine t-butyl ester (DAPT), which is known to increase the number of *Atoh1*+ secretory progenitors and decrease the number of stem cells^104,105^. Using an *Atoh1^CreERT2^:Rosa26^tdTomato^* reporter line, we treated bioprinted and manually seeded organoid cultures with DAPT for 2 or 3 days after seeding and imaged after 4 or 5 days (Extended Data Fig. 9). In manually seeded organoids, differences in overall tdTomato signal acquired by confocal microscopy were obscured by heterogeneity in organoid position, size, and shape. In contrast, there was a clear change in morphology and increase in tdTomato signal for DAPT-treated organoids in bioprinted arrays. We hypothesize this morphological change is linked to previously described impacts of Notch inhibition on intestinal stem cell proliferation, which may be more obvious in bioprinted organoid arrays (Extended Data Fig. 9)^106^.

Quantifying the volume of tdTomato-positive signal in each condition revealed that while there was a statistically significant increase in signal for both bioprinted and manually seeded organoids in treated vs. untreated conditions, the effect size was substantially improved using bioprinted arrays (∼3.7-fold increased difference between treated and untreated means) (Fig. 5h). Signal-positive volumes were also not normally distributed when manually seeded (p < 0.0001 as determined by D’Agostino & Pearson test). Bioprinted organoids, in contrast, were more normally distributed and had decreased variance (printed CoV = 48% for treated and 58% for untreated; manual CoV = 127% for treated and 174% for untreated). Computing a post-hoc power analysis (α = 0.05; β = 0.2) with the given effect sizes and variances in each condition recommended n = 12 printed organoids compared to n = 100 manually seeded organoids. This nearly order-of-magnitude decrease in required comparisons emphasizes the attractiveness of MAGIC matrix bioprinting for rare tissues or subtle phenotypes.

To more precisely quantify how bioprinted arrays improved assay sensitivity, we performed bootstrapping on the bioprinted and manually seeded populations to calculate p-value as a function of the number of paired organoid comparisons (treated vs. untreated). In the bioprinted arrays, we achieved p-values below 0.05 after comparing only 5 organoids; in manually seeded organoid cultures, 45 comparisons were required to reach the same statistical significance (Fig. 5i). P-values for bioprinted organoids continued to decrease with additional comparisons approaching p∼10^-10^ after 45 comparisons – the same number of organoids necessary to reach a p-value of 0.05 in manually seeded organoids. This 10^8^-fold decrease in p-value highlights the potential of these bioprinted organoid arrays to detect subtle phenotypes in chemical, microenvironmental, and genetic screens.

### Flexible production of 3D perfusable organoid tubes

Given the importance of fluid flow and both static and cyclic stress during processes such as peristalsis^3,107,108^, we examined whether mouse intestinal organoid tubes could be perfused following their lumenization and crypt morphogenesis (Fig. 5j). Using glass capillaries, we pierced the tissue lumen and introduced fluid using a syringe to clear internal cellular debris. By applying oscillatory fluid flow we could simulate cyclic expansion and contraction as might be experienced during peristalsis (Fig. 5k, Supplementary Movie 5). We observed radial expansion over ∼30% of the perfused tubes at peak pressure enabled by the high deformability of MAGIC matrices, and unlike cultures using materials with non-physiological stiffness such as PDMS. Thus, MAGIC matrix tube bioprinting enables access to the organoid lumen and the introduction of biophysical cues such as fluid shear and both static and cyclic stretch, key features of microphysiological systems, while maintaining a free basal surface that ultimately can be interfaced with other tissue types.

## Discussion

Promoting more reproducible and complex organoid morphogenesis requires biomaterials that mimic the dynamic boundaries encountered by tissue in vivo and that also support engineering controls necessary for setting the initial conditions of a culture. Here we developed and characterized a simple granular biomaterial that supports both powerful embedded bioprinting of cell slurries to set initial conditions and the large tissue deformations that occur during organoid morphogenesis. MAGIC matrices are highly tunable and revealed unique relationships between the loss tangent at low strains and short timescales and stress relaxation at higher strains and longer timescales. This was true for a variety of MAGIC matrix compositions, presenting a suite of 3D granular biomaterials with which to investigate rheological determinants of tissue behavior. Previous studies have demonstrated that rheological properties such as storage modulus, loss modulus, and stress relaxation are collectively important for symmetry breaking and morphogenesis in intestinal organoids but could not investigate whether stress relaxation was independently important. By preparing materials with nearly identical storage and loss moduli at short timescales, we revealed the critical importance of stress relaxation at longer timescales and larger strains, consistent with the large tissue deformations required during the morphogenesis of intestinal crypts. Another critical finding was that optimal MAGIC matrices were an order-of-magnitude softer than most synthetic matrices reported in the literature. Similarly, the yield-stress of many reported embedded printing materials is much higher than that of MAGIC matrices^62,109^, which may contribute to their ineffectiveness as cell culture materials^110^. However, the desired yield-stress of MAGIC matrices can be tuned for a given application independent of storage and loss modulus by tuning AMG:Matrigel volume fraction. MAGIC matrices are relatively simple in design, employing off-the-shelf and well-characterized constituents. Moreover, they are readily tunable by changing the size, wt.%, and volume fraction of the granular medium and interstitial material.

To take advantage of these materials in preparing more uniform and complex tissues in vitro, we developed a piezoelectric printhead that precisely aspirates and extrudes cell slurries at tissue-like densities directly into MAGIC matrices, thereby allowing tissues to autonomously self-organize while minimizing the variability typically associated with manual organoid seeding. Future work investigating more advanced bioprinter technologies such as coaxial printheads could provide access to even more advanced initial tissue-ECM architectures^111,112^. Combined, the matrix and printhead enable rapid prototyping and controlled extrusion of delicate and low-volume living materials.

To demonstrate the power of MAGIC matrices, we tuned their chemical and rheological properties to match those of rBMs (such as Matrigel) while supporting embedded 3D printing. The rheological features of rBMs that support organoid growth and morphogenesis have largely been attributed to their soft, viscoelastic nature. Our work and others further suggest that rBMs behave more like viscoelastic liquids over the length- and timescales of self-organization, and this capacity to dissipate stress may be a key driver of morphogenesis^21,113^. Synthetic biomaterials incorporating appropriate viscoelasticity and some degree of stress relaxation were also shown to support 3D symmetry breaking, spreading, invasion, and differentiation^21–23,60,67,71,114–116^. However, in all of these studies, loss tangent is strongly correlated with stress relaxation at long timescales, making it impossible to determine how stress relaxation independently contributes to tissue morphogenesis. By modulating the volume fraction of the granular medium, we tuned stress relaxation at long timescales and high strains without affecting loss tangent, allowing us to determine the quantitative impact of stress relaxation alone on crypt morphogenesis. Our findings highlight the importance of complete dissipation of internal stresses over the larger timescales and deformations relevant to morphogenesis. While the benchmark 1:1 MAGIC matrix composition with 0.5 wt% AMGs did not completely relax over the length and timescales measured, the residual stresses (∼6% or ∼0.2 Pa) were significantly below those known to be exerted by tissues during growth and morphogenesis^63,74,117^. Moreover, deformations of MAGIC matrices using nanoindentation suggest that at the tens of microns length-scale relevant to cells and tissues, MAGIC matrices may relax stresses at a rate and magnitude nearly identical to Matrigel. Among the suite of biomaterials suitable for embedded 3D tissue culture, MAGIC matrices allow for significantly greater stress relaxation at large deformations and timescales^11,17,21,47,68,71^. The tunability of stress relaxation was generalizable for the range of MAGIC matrix compositions tested, highlighting a potentially unique property of granular materials with viscous interstitium that warrants further investigation. Notably, we did not engineer the shape of the granular medium in this study, a feature predicted to control packing density^27,118^ and whose optimization may increase the range of volume fractions, and thus the degree of stress relaxation, afforded by MAGIC matrices.

Most synthetic ECMs have been designed for a particular organoid or tissue type. We took advantage of the features of MAGIC matrix that support both extrusion bioprinting and organoid culture to successfully promote the self-organization of mouse and human tissues from all major germ layers, including endoderm (intestinal), ectoderm (brain; mammary; salivary gland), and mesoderm (vasculature, fibroblasts). Focusing on intestinal organoids, we found remarkably increased homogeneity in organoid structure and phenotype by a number of metrics, including growth, morphogenesis, and maturation rate. This homogeneity yielded organoid arrays that were “assay-ready” in only 2-3 days, compared to previous studies that produced organoids of similar homogeneity but required 5 days or more in culture^14,31^. Moreover, previous reported methods for preparing organoid arrays could not simultaneously control tissue architecture beyond spheroids, nor organoid position within the embedding ECM. In addition to intestinal organoids, all tissue types could be bioprinted and cultured in high-throughput arrays or as tubes with little-to-no changes in printing parameters or matrix composition, for example including collagen or excluding Matrigel. By a variety of functional metrics, these organoids were indistinguishable from those cultured in pure Matrigel, suggesting that the unique mechanisms contributing to tissue folding, budding, and cell sorting necessary for the self-organization of these other tissue types are also supported in MAGIC matrix^65,95,119,120^.

MAGIC matrix bioprinting held other unique features that point to future applications in high-throughput biology. For example, we found that organoid arrays fully embedded in MAGIC matrix were amenable to transfection with mRNA and transduction with lentivirus far more efficiently than in Matrigel. Their uniform position facilitated both live and fixed imaging as well as automated analysis. Moreover, MAGIC matrix bioprinting resulted in dramatic improvements in assay statistical power, decreasing the predicted number of tissues required to identify a phenotype by nearly an order-of-magnitude or more and providing orders-of-magnitude decreases in confidence intervals compared to manual culture. Therefore, this platform presents opportunities to work with precious tissue sources such as primary patient biopsies in applications like clinical drug screens. Finally, these technologies provide a route to more complex and in vivo-like 3D microphysiological systems, for example by perfusing bioprinted organoid tubes. The regularity and scalability of these models could be combined with established perfusion and co-culture methods without the drawbacks of artificial interfaces and geometries. These findings lay the foundation for building even more complex and reproducible models of human and animal development, homeostasis, and disease.

## Materials and Methods

Materials and methods are provided in the supplement.

## Data and Code Availability

All source data will be made available upon peer-reviewed publication. Custom MATLAB, R, and Image J Macro (IJM) scripts will be publicly made available through GitHub upon peer-reviewed publication or can be provided upon request.

## Acknowledgements

We thank the members of the Gartner Lab, Klein Lab, and CZ Biohub Bioengineering Group for their valuable suggestions. We thank Sandra Montes-Olivas for guidance in the use of her MATLAB crypt counting software. We thank Dr. Kevin Healy of UC Berkeley for providing rheological equipment. We thank Dr. Mark Skylar-Scott of Stanford University and his group for providing technical input and assistance. This work was funded by the Chan Zuckerberg Investigator Program to Z.J.G., the UCSF Center for Cellular Construction (DBI-1548297), a CBCRP IDEA award, and the NIDDK (R01DK126376). A.J.G. was supported by a Chan Zuckerberg Biohub Collaborative Postdoctoral Fellowship. V.S. was supported by the Zena Werb Memorial Fellowship through the Helen Diller Family Cancer Center at UCSF. FACS was performed on a BD Aria 3, supported in part by HDFCCC Laboratory for Cell Analysis Shared Resource Facility through a grant from NIH (P30CA082103).

## Author Contributions

A.J.G. and Z.J.G. conceptualized the embedded bioprinting material design and application. A.J.G., M.W.L.K., H.K., P.L., R.G.S., and Z.J.G. conceptualized the bioprinter instrumentation. A.J.G. performed materials synthesis and characterization. M.W.L.K., P.L., and R.G.S. designed and built the bioprinter hardware and wrote the software. A.J.G., V.S., S.V., K.M.H, and N.G. performed primary cell isolation, culture, microscopy, immunofluorescence, and organoid experiments. A.J.G., M.W.L.K., and H.K. performed experiments to identify optimal bioprinting parameters. M.W.L.K. wrote custom scripts for bioprinting organoid arrays. A.J.G., V.S., and M.B. wrote custom scripts for microscopy and statistical analysis. K.P., C.M., and S.K. consulted on and provided experimental assistance with material characterization. V.S., S.V., K.M.H., N.G., J.M.R., N.C., T.J.N., and O.K. consulted on and provided experimental assistance with organoid isolation, characterization, and application. A.J.G. and Z.J.G. wrote the manuscript, with review and feedback from all other authors.

---

**Extended Data Figure 1.**
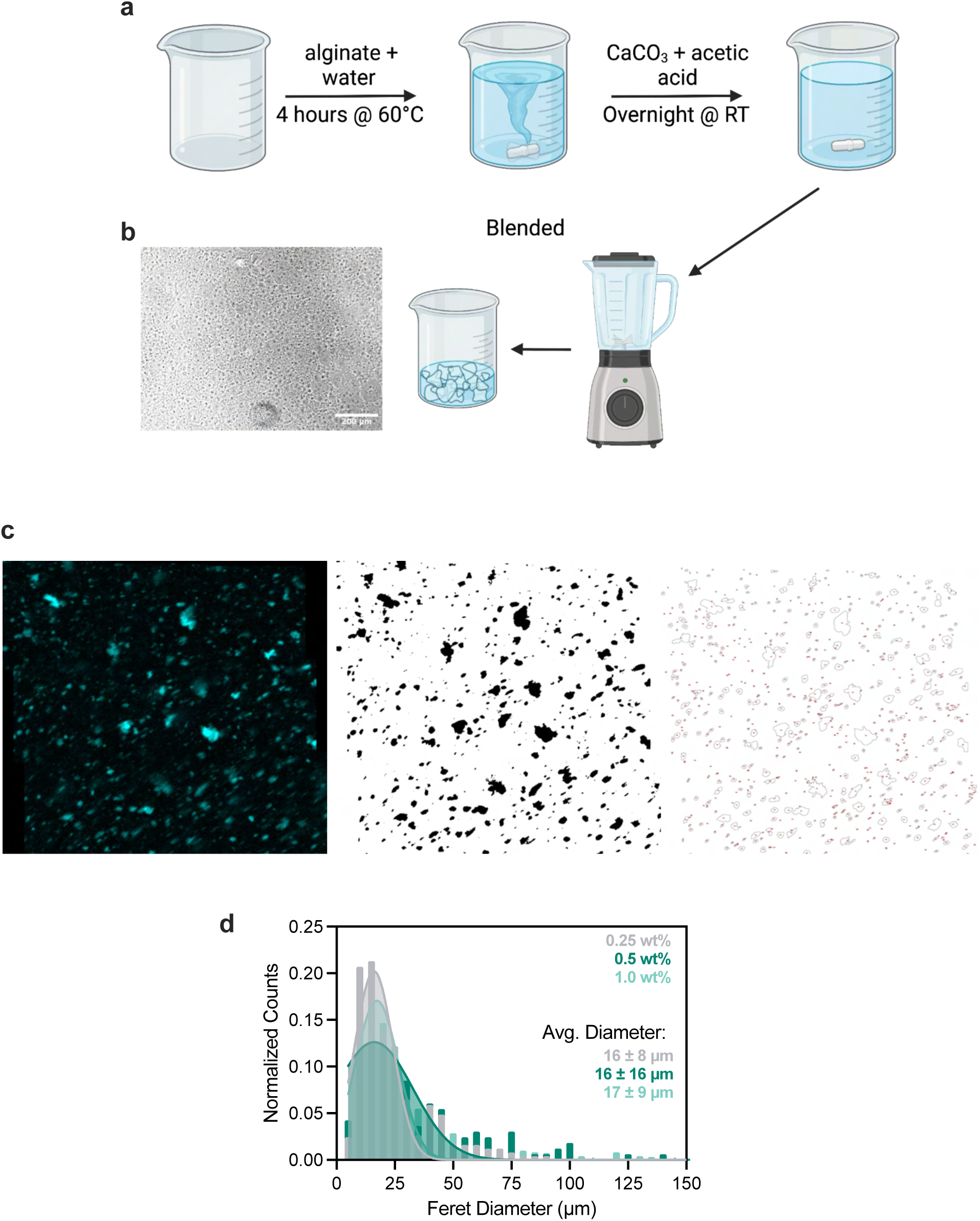
Alginate microgels are roughly cell-sized. **a.** Cartoon workflow for preparation of alginate microgel slurry. **b.** Brightfield image of microgel slurry after synthesis, with nearly transparent microgels. Scale bar = 200 µm. **c.** Representative images outlining workflow for quantifying microgel size. The polyanionic alginate backbone was positively stained with DAPI and segmented in Fiji to calculate particle diameter. **d.** Distribution of alginate microgel sizes fit to a Gaussian distribution. Data shown are mean ± SD from *n* = 3 separate images from 3 separate fields of view for each alginate wt% and 3 independent AMG preparations.

**Extended Data Figure 2.**
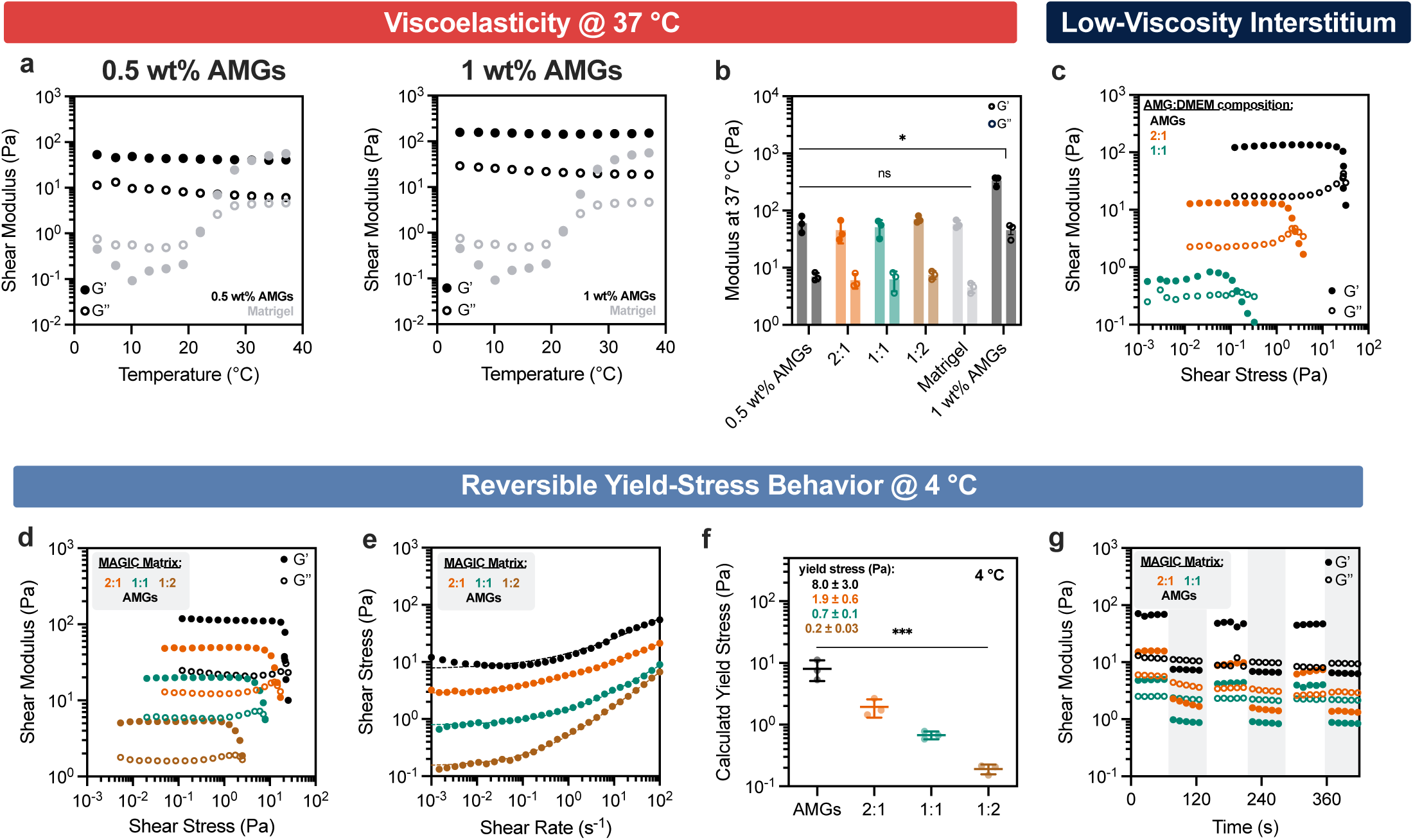
MAGIC matrices are tunable embedded printing and cell culture biomaterials. **a.** Storage and loss modulus of (left) 0.5 wt% and (right) 1 wt% alginate microgel (AMG) pelleted slurry and Matrigel as a function of temperature. **b.** Storage and loss moduli of MAGIC matrix formulations at 37 °C using pelleted 0.5 wt% AMGs or (right) 1 wt% AMG slurry at 1 Hz and 1% strain. Data shown are mean ± SD from n = 3 independent microgel preparations. * = p < 0.05 for both storage and loss modulus of all MAGIC matrix formulations and 1 wt% AMG slurry compared to pure Matrigel as determined by one-way ANOVA with Dunnett’s multiple comparisons. **c.** Oscillatory amplitude sweeps at 4 °C for various compositions of pelleted AMGs or AMGs diluted in cell culture media as a low-viscosity interstitium compared to Matrigel, which decreases jamming and yield-stress. 1:2 AMG:DMEM compositions did not exhibit yielding behavior. **d.** Oscillatory amplitude sweeps at 4 °C for various MAGIC matrix compositions show yielding behavior indicated by G’ and G’’ cross-over. Data shown are representative of n = 3 independent microgel preparations. **e.** Unidirectional shear rate measurements at 4 °C fit to a Herschel-Bulkley power law model. **f.** MAGIC matrix yield stress values calculated using Herschel-Bulkley fits in (D). Data shown are mean ± SD of n = 3 independent microgel preparations. **g.** Reversible yield-stress test wherein applied strain is alternated between 1% and 100% for a variety of MAGIC matrix formulations at 4 °C. Gray bars indicate areas of 100% strain. Cross-over and recovery of G’ and G’’ indicates reversible viscoelastic behavior. For **d**, **e**, and **g**, data are representative of *n* = 3 independent microgel preparations.

**Extended Data Figure 3.**
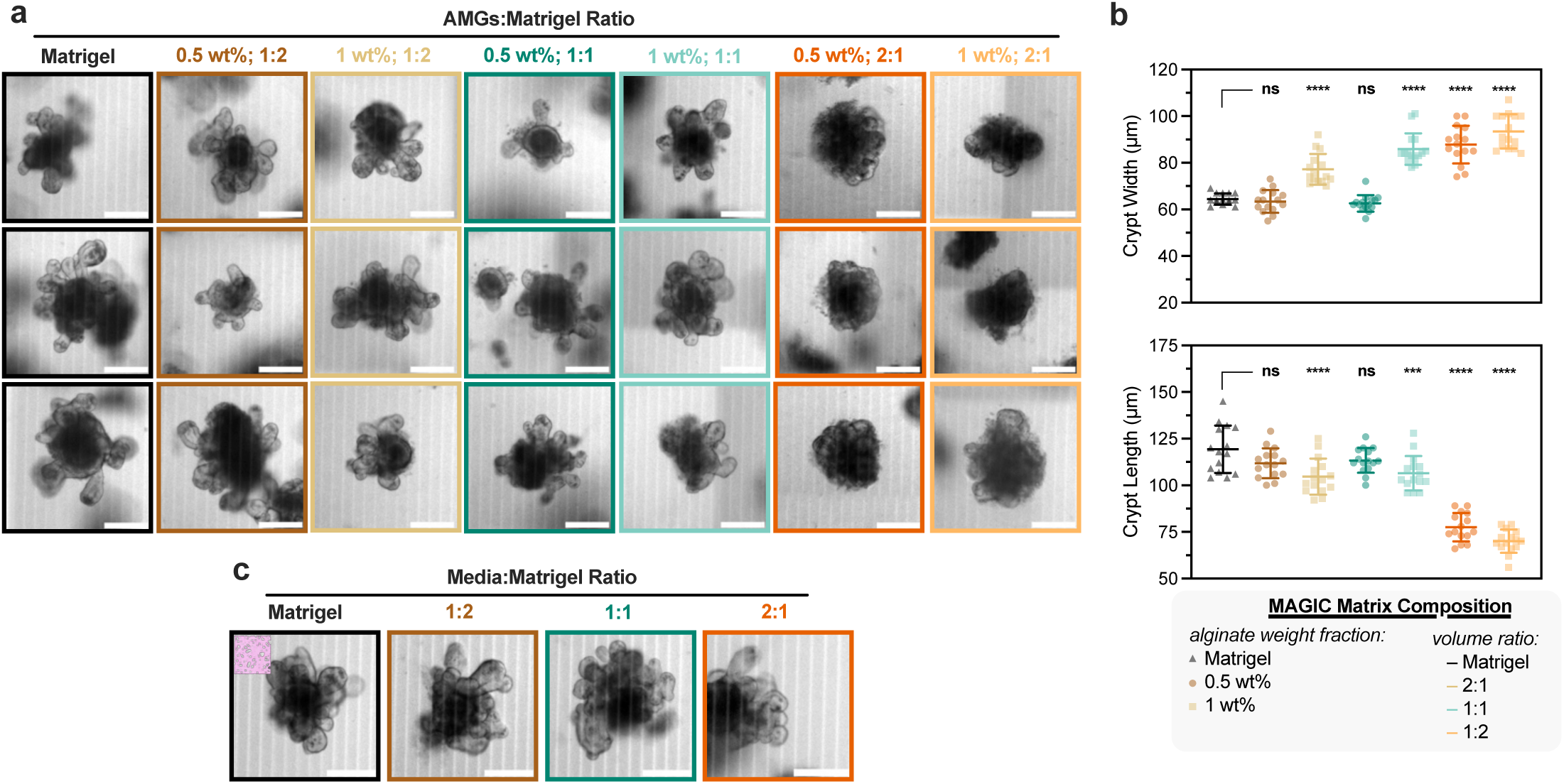
MAGIC matrix composition impacts organoid morphogenesis. **a.** Representative images of mouse intestinal organoids at 5 days after manually seeding into Matrigel and MAGIC matrices of varying AMG wt% and composition with Matrigel. MAGIC matrix compositions are represented as added volume ratio of AMGs:Matrigel. Scale bars = 200 µm. **b.** Quantification of organoid crypt width (top) and crypt length (bottom) as a function of matrix composition. **c.** Representative images of organoids grown in Matrigel diluted at various volume ratios with mouse intestinal organoid growth medium. Scale bars = 200 µm.

**Extended Data Figure 4.**
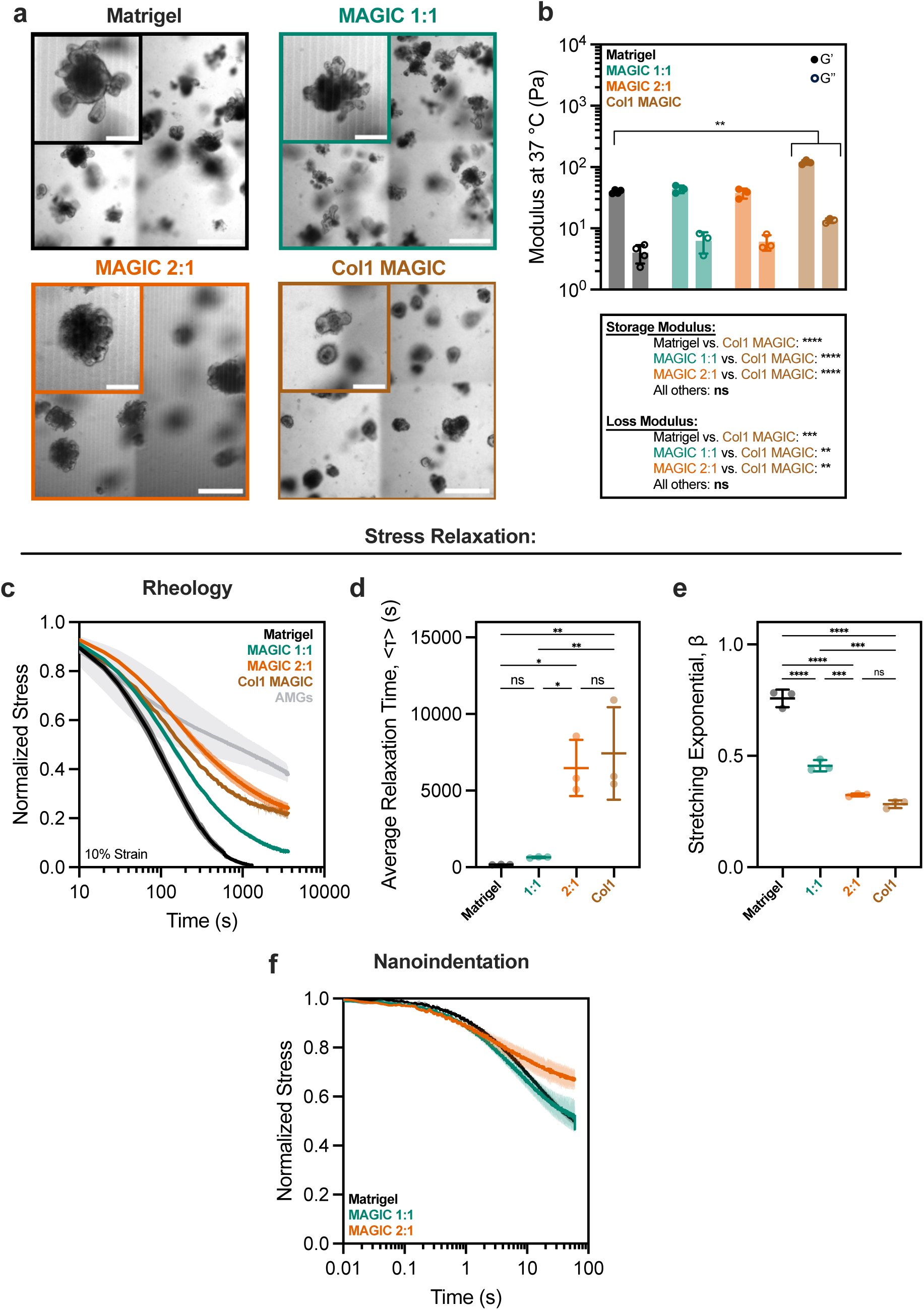
ECM stiffness and stress relaxation determine phenotypes of crypt morphogenesis in intestinal organoids. **a.** Representative images of mouse intestinal organoids at 5 days after manually seeding into Matrigel, MAGIC matrices, and collagen 1-containing matrices. Collagen 1-containing matrices were composed of 25% 4 mg/mL Bovine Col-1, 25% Matrigel, and 50% pure AMGs by added volume (final concentration of 1 mg/mL Col-1). Scale bars = 500 µm. Insets show zoomed in view of representative organoids. Scale bars = 200 µm. **b.** Storage and loss moduli of Matrigel, MAGIC matrix, and Collagen I-containing matrices at 37 °C, 1 Hz, and 1% strain. Table summarizes statistical analysis. **c.** Normalized stress relaxation curves for Matrigel, AMGs, MAGIC matrices, and Collagen 1-containing matrices over 1 h at 10% strain. **d, e.** Quantification of average relaxation time (**d**) and stretching exponential € for each matrix determined using a stretched exponential model. **f.** Normalized stress relaxation measurements of Matrigel and MAGIC matrices of various compositions using nanoindentation at 50 µm indentation for 60 s. Data shown are mean ± SD from n = 3 independent ECM preparations. * = p ≤ 0.05; ** = p ≤ 0.01; *** = p ≤ 0.0001; **** = p ≤ 0.00001; ns = not significant as determined by one-way ANOVA with Tukey’s multiple comparisons.

**Extended Data Figure 5.**
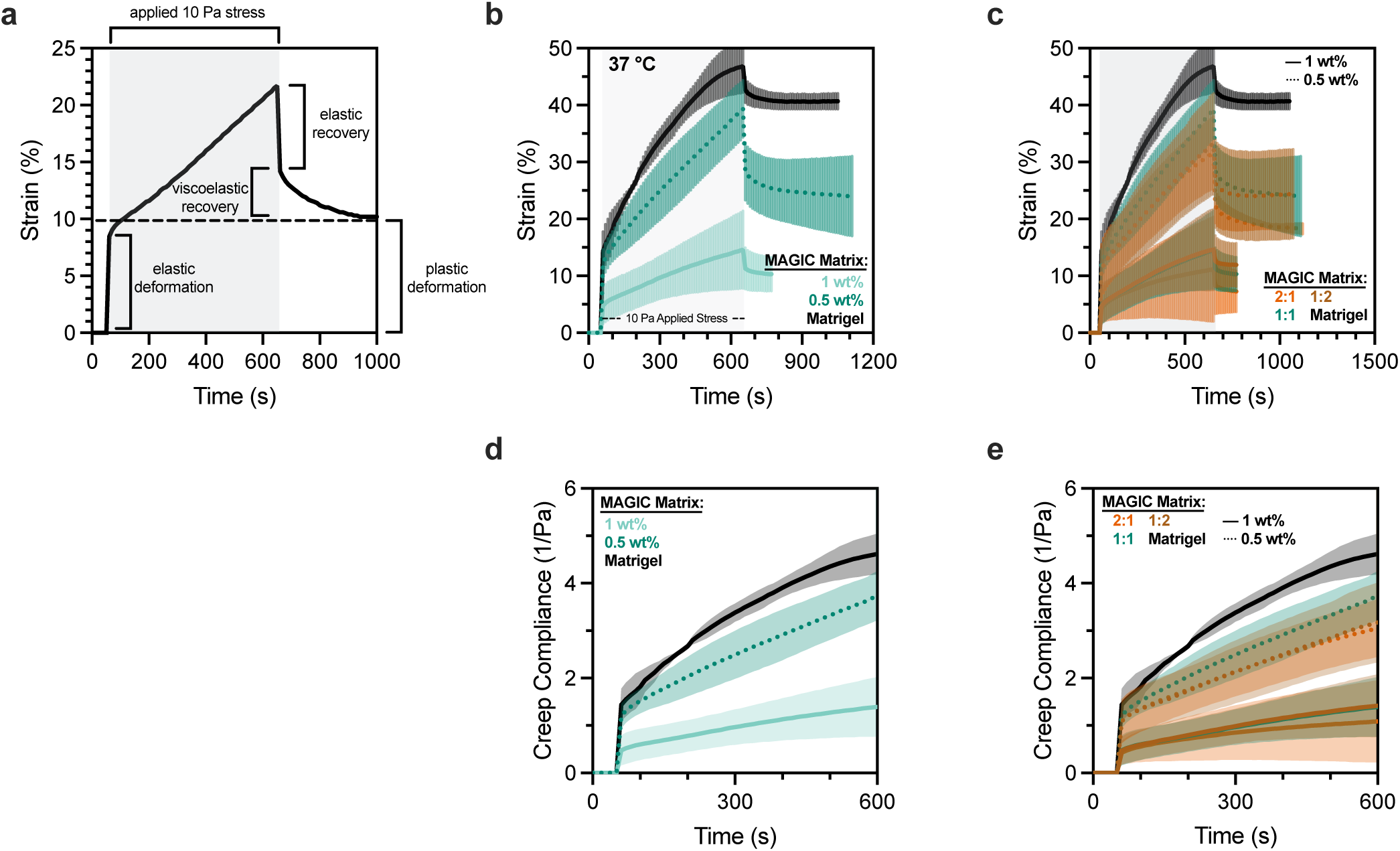
MAGIC matrices exhibit composition-dependent plasticity and creep. **a.** Representative data illustrating different mechanical modes of a creep-recovery test meant to simulate tissue expansion with constant force. **b, c.** Creep-recovery test for 10 Pa applied constant shear stress (gray bar) for different MAGIC matrix formulations measuring material strain rate compared to pure Matrigel. **d–e.** Creep compliance curves measured during creep experiments shows corresponding differences in matrix relaxation. In general, response to applied stress is a strong function of alginate wt%, but not fraction of Matrigel. Data shown are mean ± SD from n = 3 independent microgel preparations.

**Extended Data Figure 6.**
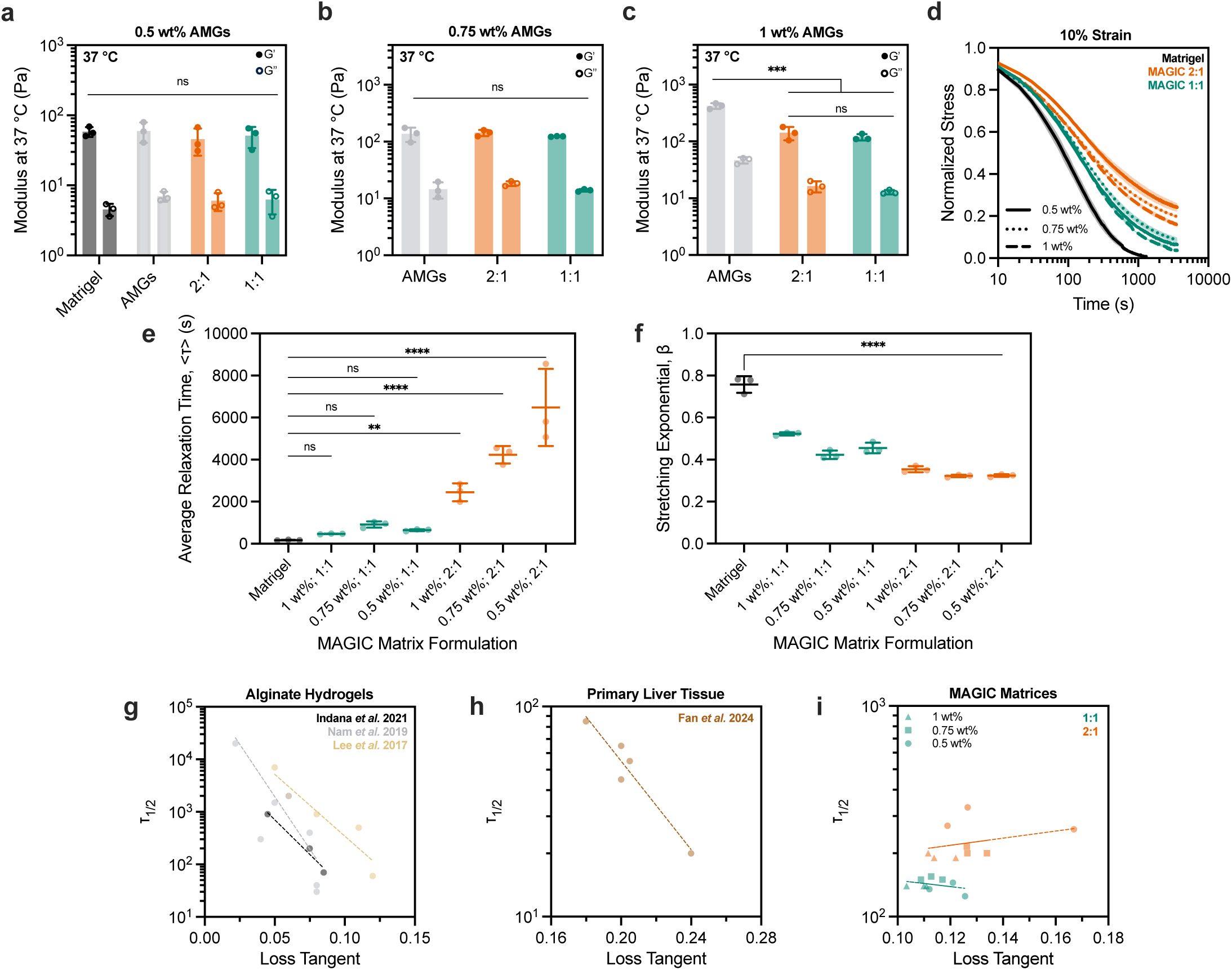
MAGIC matrices of various compositions tune stress relaxation independent of storage and loss moduli. a–c. Storage and loss moduli of MAGIC matrix formulations at 37 °C using 0.5 wt% (**a**), 0.75 wt% (**b**), or 1 wt% (**c**) AMGs at 1 Hz and 1% strain. Data shown are mean ± SD from n = 3 independent microgel preparations. **d.** Normalized stress relaxation curves for Matrigel or MAGIC matrices over 1 h at 10% strain. **e–f.** Quantification of average relaxation time (**e**) and stretching exponential (**f**) for MAGIC matrix formulations using a stretched exponential function. **g–i.** Time at half-maximum stress relaxation, or τ1/2, compared to loss tangent showing negative correlation for a variety of alginate hydrogels (Indana *et al*. 2021, R^2^ = 0.99; Nam et al. 2019, R^2^ = 0.80; Lee et al. 2017, R^2^ = 0.63) and primary liver tissue (Fan *et al.* 2024, R^2^ = 0.88) from published datasets, but not for MAGIC matrices (1:1, R^2^ = 0.11; 2:1, R^2^ = 0.09). Linear fits are shown as a guide-to-the-eye. * = p ≤ 0.05; ** = p ≤ 0.01; *** = p ≤ 0.0001; **** = p ≤ 0.00001; ns = not significant as determined by two-way ANOVA with Tukey’s multiple comparisons (**a–c**) or one-way ANOVA with Dunnett’s multiple comparisons (**e–f**).

**Extended Data Figure 7.**
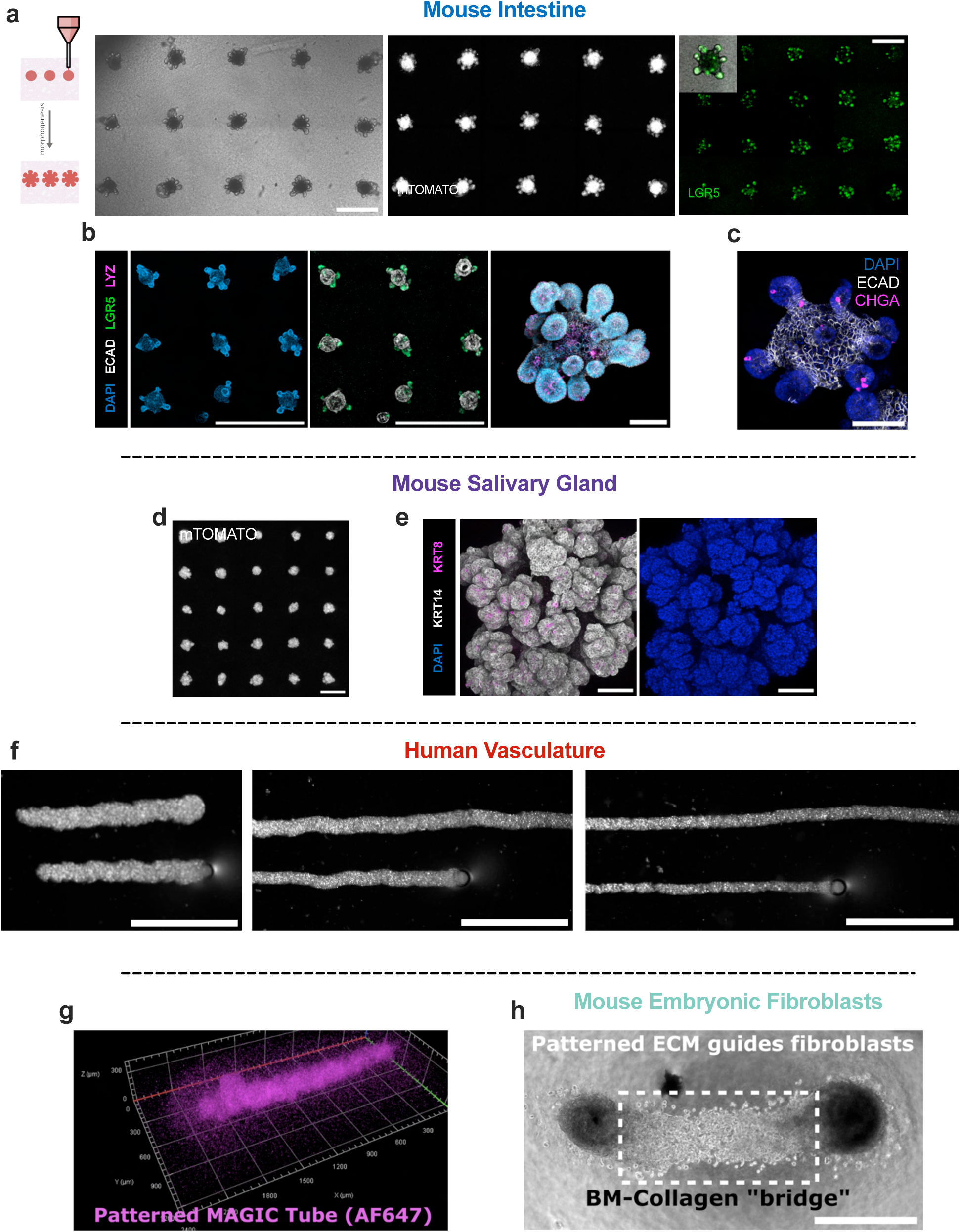
MAGIC matrices promote canonical self-organization of various organoid types from different developmental lineages. **a.** Brightfield and fluorescent live images of mouse intestinal organoid arrays 2 days after bioprinting expressing either mTomato or *Lgr5*-eGFP. eGFP signal localizes to the base of the crypts, where *Lgr5*+ stem cells should reside. Scale bars = 500 µm. **b.** Left, maximum intensity projections of DAPI, *Lgr5*-GFP, ECAD-stained intestinal organoid arrays 3 days after bioprinting. Scale bar = 1 mm. Right, 3D rendering of fixed bioprinted intestinal organoid stained for DAPI, ECAD, and Paneth cells (LYZ). Scale bar = 200 µm. **c.** Maximum intensity projection of DAPI, ECAD, and enteroendocrine cells (CHGA). Crypts protrude in all directions, highlighting fully 3D morphogenesis in MAGIC matrices. Scale bar = 200 µm. **d.** Live imaging of bioprinted salivary gland organoid arrays expressing mTomato. Scale bar = 1 mm. **e.** Immunofluorescence of fixed bioprinted salivary gland organoids showing presence of both basal (keratin 14) and ductal (keratin 8) cells. Scale bar = 100 µm. **f.** Brightfield images of HUVEC cords bioprinted at different stage translation speeds highlighting control over vessel width. Scale bars = 1 mm. **g.** 3D rendering of an AF647-NHS stained MAGIC matrix tube printed into an AMG bath, highlighting the ability to pattern ECM. **h.** Brightfield images of mouse embryonic fibroblasts printed as two nodes with patterned Collagen 1 and MAGIC matrix printed between the nodes. Fibroblasts spread only into patterned area, highlighting spatial control over cell migration. Scale bar = 500 µm.

**Extended Data Figure 8.**
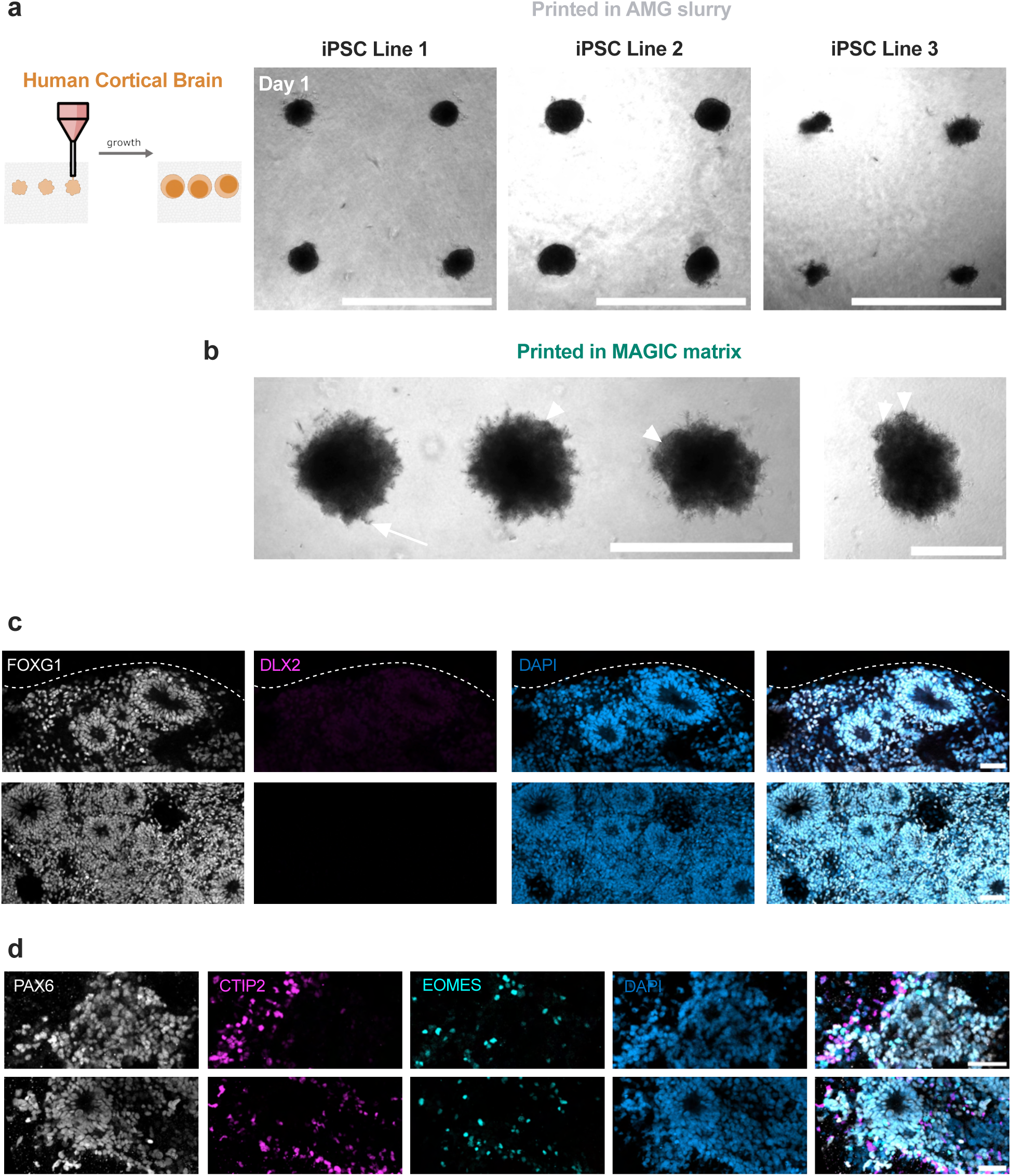
Bioprinted hiPSC-derived cortical brain organoids exhibit matrix-dependent phenotypes and rosette organization. **a.** Live images of human iPSC-derived cortical organoids from three different donors (methods) bioprinted into arrays using AMG support baths. Scale bar = 1 mm. **b.** Cortical organoids bioprinted in MAGIC matrices show sprouting (arrow) and neuroepithelial budding (arrowheads). Scale bars = 1 mm (left) and 500 µm (right). **c, d.** 20 µm maximum intensity projections of bioprinted cortical organoids stained for (**c**) cortical identity and (**d**) neuronal differentiation. Scale bars = 50 µm.

**Extended Data Figure 9.**
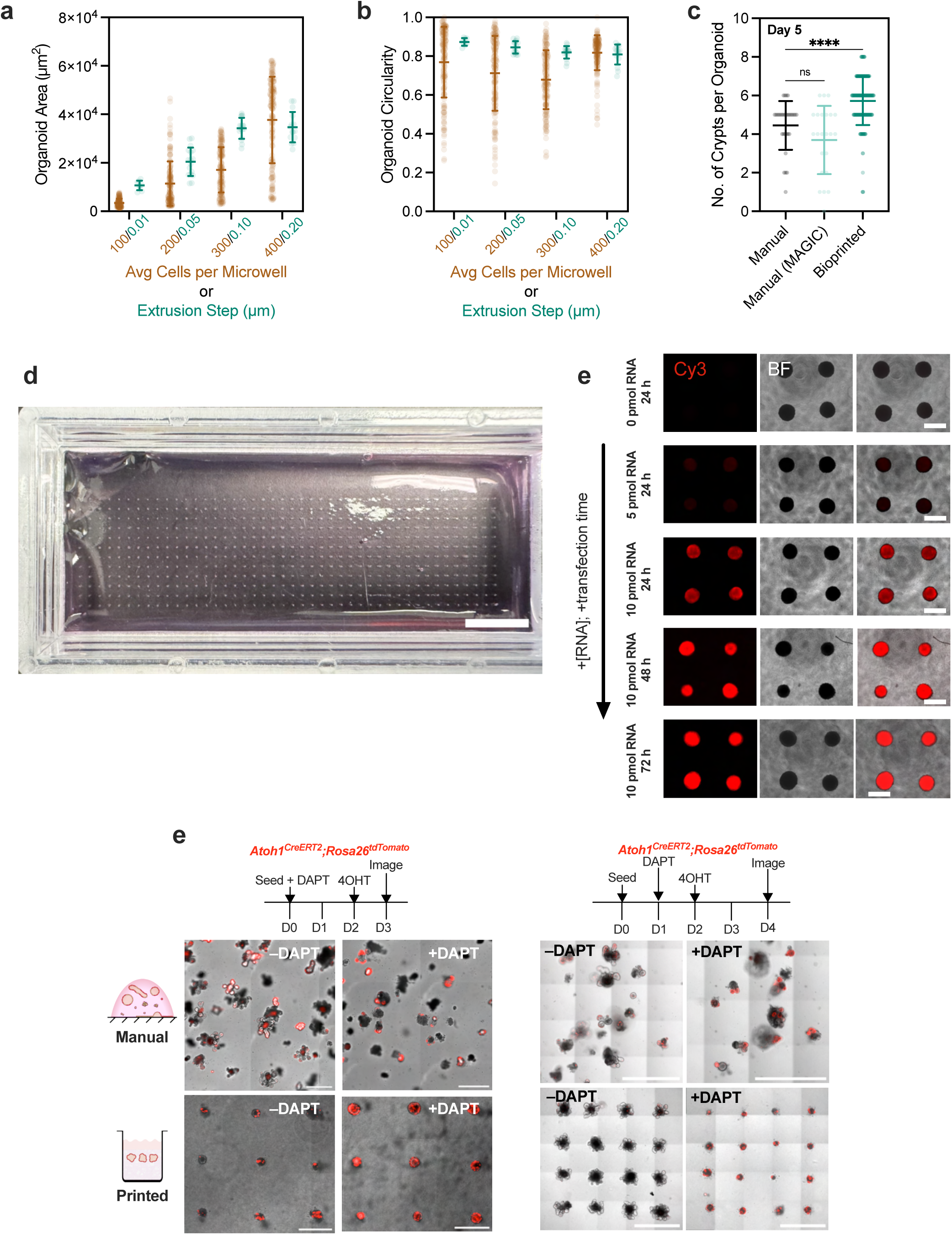
Bioprinted organoid arrays exhibit improved homogeneity and morphogenesis. **a.** Organoid area and **b.** circularity measured using max intensity projections of confocal z-stack images of GFP-expressing Caco-2 cell slurries immediately after seeding in microwells or bioprinting. **c.** Picture of large (n = 528) organoid array bioprinted into a chambered slide using MAGIC matrix. Scale bar = 10 mm. **d.** Quantification of crypts per organoid for those manually seeded in Matrigel or MAGIC matrix, or bioprinted. Data shown are mean ± SD of n ≥ 30 organoids; ns = not significant; **** = p < 0.0001 determined by one-way ANOVA with Dunnett’s multiple comparisons. **e.** Live images of bioprinted TNBC organoid arrays transfected using Lipofectamine RNAiMAX and a Cy3-conjugated, single-stranded non-coding small RNA (36mer) at various concentrations and transfection times. Scale bar = 500 µm. **f.** Live images of manually seeded and bioprinted *Atoh1*-tdTomato organoid arrays using two different DAPT treatment timelines. The left treatment timeline and images were used for bootstrapping and statistical analysis.

**Supplementary Movie 1. Dense cell slurry bioinks printed into pure Matrigel at 4 °C do not retain their printed shape.** Scale bar = 1 mm.

**Supplementary Movie 2. Example of high-throughput mouse intestinal organoid array generation using MAGIC matrix bioprinting.** Video is displayed at 8x real-time. Scale bar = 1 mm.

**Supplementary Movie 3. MAGIC matrix bioprinting of HUVEC vascular cords of different widths.**

**Supplementary Movie 4. MAGIC matrix bioprinting supports patterning of centimeter-scale tissues.** Scale bar = 1 mm.

**Supplementary Movie 5. Cyclic 3D pressurization of a perfused intestinal organoid tube.** Tube diameter increases and decreases upon application and removal of pressure, indicating the tissue is experiencing strain.

