## Supplementary Information for "Stress relaxing granular bioprinting materials enable complex and uniform organoid self-organization"

[1]Department of Pharmaceutical Chemistry, University of California San Francisco, San Francisco, CA
[2]Chan Zuckerberg Biohub SF, San Francisco, CA
[3]Department of Orofacial Sciences, University of California San Francisco, San Francisco, CA
[4]Department of Bioengineering, University of California Berkeley, Berkeley, CA
[5]Weill Institute for Neurosciences, University of California San Francisco, San Francisco, CA
[6]Department of Medicine, University of California San Francisco, San Francisco, CA
[7]Graduate Program in Bioengineering, University of California San Francisco, San Francisco, CA and University of California Berkeley, Berkeley, CA
[8]Department of Chemical and Biomolecular Engineering, University of California Berkeley, Berkeley, CA
[9]Department of Bioengineering and Therapeutic Sciences, University of California San Francisco, San Francisco, CA
[10]Department of Mechanical Engineering, Stanford University, Stanford, CA
[11]Department of Bioengineering, Stanford University, Stanford, CA
[12]Department of Pediatrics, Cedars-Sinai Guerin Children's, Los Angeles, CA
[13]Center for Cellular Construction, University of California San Francisco, San Francisco, CA

### 1. Stress Relaxation Quantification

To quantify the relevant time scale of stress relaxation we fit stress relaxation profiles using a stretched exponential function[1] :

;

where σ describes shear stress over time, β is the stretching exponent, and $\tau_c$ is the characteristic relaxation time. Values of β for Matrigel were closer to 1, indicating that the stress response is dominated by a single relaxation mode. Values of 1:1 and 1:2 MAGIC matrix lie around 0.5 and 0.3, respectively (Extended Data Fig. 3). These results suggest that the relaxation modes in MAGIC matrices are more complex as the microgel fraction increases, which appears to impact long-term tissue behavior. We additionally quantified an average relaxation time for each material as recently described[1]:

.

For comparison of loss tangent and stress relaxation, the stress relaxation half-life, or $\tau_{1/2}$, was calculated from stress relaxation curves or obtained from published datasets[2–5] and plotted as a function of loss tangent for each reported material formulation. Data were fit using linear regression to demonstrate negative correlation between loss tangent and stress relaxation half-life for all analyzed materials besides MAGIC matrices. Data were plotted based on material identity, including alginate-based materials, primary liver tissue, and MAGIC matrices to account for material-specific differences in half-life magnitude.

### 2. Materials & Methods

*1.1 MAGIC matrix preparation*: Sodium alginate (Sigma, 9005-38-3) is dissolved at 1 wt% (typically 1 g into 100 mL) into pre-heated 60 °C sterilized $ddH_2O$ and stirred for 2–4 h until the solution is homogeneous. In a separate flask, calcium carbonate is mixed at 0.2 wt% (typically 200 mg into 100 mL). The dispersed calcium carbonate suspension is added to the alginate solution and cooled to room temperature while stirring for 1 h, leading to 0.5 wt% alginate and 0.1 wt% $CaCO_3$. Pure acetic acid (Sigma, 64-19-7) is added in drop-wise, slowly at a 1:500 ratio (typically 400 µL) under constant stirring at 1000 RPM. The solution should increase in viscosity, indicating release of $Ca^{2+}$ ions and alginate cross-linking. The solution is left stirring overnight at 1000 RPM to shear the mixture, leading to microgel formation. The next day, this mixture is blended for 60 s on the High setting using a Hamilton Beach commercial blender (BioSpec 908). The microgel mixture is then strained through a 100 μm filter to remove large particles. The filtered microgels are centrifuged at 20,000xg for 20 min at 4 °C and the supernatant aspirated, followed by resuspension in 1x volume of DMEM:F-12 (UCSF Cell Culture Facility) medium supplemented with penn/strep, Primocin (Invivogen), and 2 mM NaOH, and stored at 4 °C overnight or until the day before use. The day before use, the mixture is again centrifuged under the same conditions and the media replaced with 2x volume DMEM:F-12 supplemented with penn/strep, Primocin, 4% v/v HEPES (1 M, UCSF Cell Culture Facility), and 4% v/v $NaHCO_3$ (37 g/L in water, pH = 9.5 with NaOH). This mixture is left overnight, centrifuged the next day, and the supernatant aspirated. The resulting packed microgels are considered an undiluted microgel slurry and are stored at 4 °C until use.

Just before seeding or printing, the undiluted slurry was mixed in various ratios (typically 1:1 pipetted volume ratio) with liquid basement membrane (typically Growth Factor-Reduced Matrigel, Corning 354230) at 4 °C. This leads to the desired dilution of microgels in basement membrane, or MAGIC matrix, for seeding or printing. The matrix is kept cold throughout seeding or printing, and then moved to a $CO_2$ incubator at 37 °C for 5-10 min after seeding or printing to allow for basement membrane cross-linking. Cell culture media specific to the bioprinted tissue type is then added.

*1.2 Rheological measurements and analysis:* All rheological measurements were performed using a 25 mm cone-and-plate geometry on an Anton-Parr rheometer. For shear oscillatory measurements, both frequency and amplitude sweeps were performed, with constant 1% strain and 1 Hz frequency, respectively. Temperature sweeps were also performed at 1% strain and 1 Hz frequency, with 60 s for temperature equilibration before each measurement. For reversible shear experiments, 1% or 100% strain was applied at 1 Hz for 1 min each. To calculate the specific yield-stress of each composition, unidirectional shear measurements were performed at 1% strain and the data were fit to a Herschel-Bulkley Power Law model. The y-intercept was used to determine the yield-stress. For stress relaxation experiments, the same cone-and-plate geometry was used to apply 1, 10, or 100% strain over 1 h after cross-linking by temperature ramp for 5 min and sealing the hydrogels with mineral oil to prevent evaporation. Nanoindentation experiments were performed using an Optics11 Life Chiaro Nanoindenter with a 50 μm tip and 0.5 N/m cantilever on indentation mode with 5 μm indentation for 60 s.

*1.3 Intestinal and salivary gland organoid isolation:* Intestinal organoids were generated from the duodenum of $Lgr5^{DTR}$ mice[6] using previously described protocols[7]. Briefly, the proximal portion of the small intestine was isolated, filleted open, and rinsed 2x in PBS + Pen/Strep on ice. The tissue was then incubated in Harvest Buffer (PBS + Pen/Strep + 2 mM DTT + 10 μM Y-27632+ 1 mM EDTA) with gentle rocking on ice for 15 minutes. After 15 minutes, the tissue was shaken for 1 minute, transferred to Crypt Dissociation Buffer (PBS + Pen/Strep + 2 mM DTT + 10 μM Y-27632+ 5 mM EDTA), and incubated for 1 hour with gentle rocking on ice. The tissue was then shaken for 1 minute, filtered through a 70 μm filter, rinsed with 5 mL of ice cold Basal Medium (Advanced DMEM/F12 High Glucose, N-acetyl-L-cysteine, Pen/Strep, Glutamax, HEPES) and enumerated. Approximately 1000 crypts were plated per well in preheated 24-well plates in 50 μL Matrigel domes (356231, Corning) in ENR medium (Basal Medium, B27 Supplement, N2 Supplement, hrEGF (50 ng/mL), Noggin (100 ng/mL), and R-spondin conditioned medium).

Salivary gland organoids were generated from the submandibular glands (SMGs) of adult female mice (Krt14-Cre; mTmG). SMGs were dissected out and washed in ice cold PBS. SMG tissue was minced into small pieces with surgical scissors, followed by thorough mincing with a razor blade. The resulting tissue fragments were resuspended in Disaggregation Medium consisting of Serum Free Medium (RPMI1640, 1% Pen/Strep, Sodium Pyruvate, MEM non-essential amino acids, L-glutamine and HEPES) with DNAse (100 μg/mL) and Liberase TL (125 μg/mL). Tissue was incubated with shaking for 1 hour at 37°C. At 30 and 60 minutes, tissue was triturated to promote dissociation. Disaggregated tissue was filtered through a 70 μm filter, washed with 3 mL of Basal Medium and enumerated. Approximately 10,000 tissue fragments were plated per in 50 μL Matrigel domes (356231, Corning) in ENR medium supplemented with Y27632 (10 μM), Fgf10 (100 ng/ml) and Fgf2 (25 ng/ml).

*1.4 Intestinal cell and organoid culture:* In general, cells or organoids were cultured according to standard protocols specific to that cell type. Caco-2 cells were cultured in DMEM + 10% FBS with media changes every 3–4 days and passaged once per week. For mouse proximal small intestinal organoids (gut organoids),

organoids were cultured according to published protocols[6–8]. Briefly, gut organoids were cultured in 3D Matrigel domes on 24-well plates with 1 mL of ENR medium (EGF, Noggin, R-Spondin) for 4–7 days before passaging, with media changes every 2–4 days, depending on desired growth rate. For passaging, gut organoids were collected by mechanical dissociation of the Matrigel domes using 1 mL cold basal medium (BM) and centrifuging. All centrifugation steps were performed at 160 xg and 4 °C for 4 min unless stated otherwise. The supernatant was aspirated, and the pellet was fragmented by pipetting up and down in 1 mL BM using a non-filtered 10–100 µL pipette tip attached to the end of a 1000 µL pipette tip to gently increase shear. The crypt fragments were collected by again centrifuging and the supernatant was aspirated. These fragments were resuspended in Matrigel, generally at a 1:4 or 1:6 split ratio (e.g. 100 or 150 µL of Matrigel for one 25 µL dome), and plated onto the bottom of 24-well plates. The plate was then inverted to prevent organoid settling, and the Matrigel allowed to polymerize in a $CO_2$ incubator at 37 °C for 5-10 min before adding 1 mL of warm ENR to each well. Gut organoids that were to be used for bioprinting were instead cultured in 1 mL ENRCV (with added 3 µM CHIR99021 and 1 mM valproic acid) to promote stem cell expansion. Salivary gland organoids were cultured using a similar method and medium as described above, using ENR supplemented with Y27632 (10 µM), Fgf10 (100 ng/mL), and Fgf2 (25 ng/mL) during maintenance as well as before printing. These organoids were also collected, dissociated, and bioprinted using the same method as the small intestine organoids.

*1.5 MAGIC Matrix Bioprinting Workflow*: Mouse intestinal organoids were collected for bioprinting 2–3 days after passaging, when the culture consists mostly of large clear cysts without significant internal dead cell debris, and prepared for single-cell dissociation as described previously[9,10] . For most bioprinting experiments, either 10 or 20 Matrigel domes were collected, yielding roughly 1–2 million cells. As during passaging, gut organoids were collected by mechanical dissociation of the Matrigel domes. After centrifugation, the pellet was gently resuspended in dissociation medium consisting of TrypLE Express (Gibco) supplemented with 2000 U•mL$^{-1}$ DNase I (STEMCELL Technologies), 1 mM N-acetylcysteine (Sigma), and 10 µM Y-27632 (R&D Systems). 1 mL of this mixture was used for every ~5 wells used for printing, so generally 2 or 4 mL. The resuspended cells were incubated for 10 min in a water bath at 37 °C, with gentle shaking to agitate the pellet every 5 min. This mixture was neutralized using 8 mL or 16 mL BM supplemented with 10% FBS and gently pipetting up and down. Using a serological pipette, the cells were extruded drop-wise through a 40 µm strainer to ensure a roughly single-cell suspension and filter debris. This mixture was centrifuged and the supernatant aspirated. The pellet was resuspended in 1 mL ENR supplemented with 2.5 µM thiazovivin (Stemgent) and 2 mM EDTA (Gibco) and centrifuged again. This pellet was resuspended in ~30 µL of ENR with thiazovivin and EDTA and transferred to the 384-well collection plate on the bioprinter. Caco-2 cell slurries and salivary gland organoid slurries were prepared using the same protocol.

As described above, the undiluted slurry was mixed in a 1:1 pipetted volume ratio with liquid reconstituted basement membrane matrix (typically Growth Factor-Reduced Matrigel, Corning 354230) at 4 °C to create MAGIC matrix. 100 µL of MAGIC matrix was then deposited into each well of a chilled 96-well plate (printbed plate) that was expected to receive cell slurry using a positive-displacement pipette to minimize bubbles. The printhead was then equipped with a 75, 125, or 200 µm ID plastic denudation micropipette (CooperSurgical, EZ-Tip) to use as the print nozzle. For most experiments, a 125 µm ID tip was used. The 384-well bioink plate was centrifuged at 100xg for 1 min to pellet the cell slurry bioink at the bottom of the well. This slurry was loaded via direct aspiration (of ≤0.66 µL) directly into the printhead nozzle to reduce dead volume and minimize required cell slurry volume for printing. The motorized stage and print dimensions were calibrated manually and print parameters set using a custom MATLAB script controlling both the microscope stage and the printhead. This allowed for printing of spheroids or tubes in custom arrays with defined inter-organoid spacing, number of organoids in an array, depth of printed organoid in the matrix, and extruded volume of cell slurry at 4 °C into MAGIC matrices. Most frequently, 3x3 or 4x4 arrays of ~200 µm-wide organoid spheroids were deposited with 750 or 1000 µm inter-organoid spacing.

After printing, the printbed plate was carefully moved to a 37 °C $CO_2$ incubator and allowed to sit for 5-10 minutes to allow for basement membrane cross-linking. 200 µL of warm ENR media with 2.5 µM thiazovivin was added to each well. Media was changed every 2–4 days for spheroids, every 2 days for tubes.

*1.6 Piezoelectric Extrusion Bioprinter Design & Operation:* The piezoelectric printhead was mounted at a fixed XY-position on a cantilevered arm fastened to a Zaber LRQ075 stage for motorized Z-control. The Leica microscope's DMI8 stage controller held the print plate and provided micron-scale control of XY-position and the resulting print shape. Microscope integration provided real-time imaging during the printing process, which allowed rapid identification and diagnosis of printing issues should they occur. The bioink plate holder was mounted to two Zaber LSQ150 stages for motorized access to each bioink stored in a 384-well sample plate. Bioinks were loaded by directly aspirating a user-specified volume from a well so that only the printed volume was loaded – this is especially beneficial for precious samples such as biopsies or cell populations with insufficient bioink volume for loading typical commercial bioprinting syringes. Aspiration and extrusion was driven by Physik Instrumente's PI-841.10 piezo, which was mounted against a fluid-filled cavity with a polyether ether ketone (PEEK) diaphragm at the interface. A solenoid valve toggled fluidic connection of the printhead cavity to a fluid reservoir, allowing the printhead to be filled, cleaned, or purged, and then sealed for single-ended printing operation. This equipment results in a volume displacement resolution of ~10 pL, a maximum aspiration/extrusion volume of ~660 nL, and a maximum theoretical aspiration/extrusion rate of ~300 μL/s (the actual rate will be highly dependent on the bioink rheological properties). The entire process, including imaging, motion control, piezo-pipetting, and fluidics, was controlled using MATLAB. Custom printing protocols were scripted for maximum repeatability, efficiency, and iterative troubleshooting (Supplementary Movie 2).

The printbed plate was chilled during the experiment using cold, dry compressed air to prevent MAGIC matrix cross-linking during printing without obscuring or wetting the optics. The printbed plate temperature was monitored using a thermocouple and kept at 4–8 °C. The bioink plate was cooled with a closed-loop recirculating chiller held at 5 °C. Before each printing experiment, a calibration procedure was completed. First, to passivate and increase hydrophilicity of the inner tubing, cavity, and pipette tip surfaces, the system was incubated for 10 minutes with a treatment solution consisting of phosphate-buffered saline (PBS), 10 mg/mL bovine serum albumin (BSA), 10 mg/mL Tetronic 90R4 (Sigma 435546 functional oligomer), and 5 mM EDTA to reduce adherence of cellular components and Matrigel. It also reduced adherence of air bubbles, which increase the system's hydraulic compliance and compromise pipetting precision and responsivity. Next, the pipette tip was centered within the field of view using the manual centering screws on a Thorlabs CXY1 stage. The bioink plate wells were then centered relative to the pipette tip using a three-corner calibration scheme. Finally, with the desired ink well in place, the desired aspiration height of the printhead pipette tip was set, typically just above the bottom of the bioink well in order to collect dense cell slurry.

Once the instrument was calibrated, a series of scripts executed the printing procedure according to user-specified parameters. These parameters were defined in an editable file of constants, and included specifications such as print shape and size, array formatting, as well as rate and amount of bioink that was loaded. The first script prompted the user to define the print area by selecting four corners of a bounding box within the print well. The printed array or tubes were constrained by the bounding box so that the tip never collided with the well walls during printing. Next, the slurry was loaded from the ink well to the tip. The tip navigated to its calibrated loading height, then aspirated slurry according to the user-specified parameters. Next, the tip navigated to the bounding box and started printing the array or tubes. The array positions were computed according to user-specified row and column dimensions while staying within the bounding box. A small back-pressure was applied by changing voltage applied to the piezo, immediately followed by a rapid z-axis, and sometimes followed by an xy-axis offset, to detach the cell slurry from the tip after each bolus. In general, a negative extrusion step of 0.01 μm and z-axis translation of 100 μm at 1 mm/s consistently detached the slurry with minimal disturbance to the printed bolus or tube. This process was repeated for each array element within a print well and could be programmed to repeat across multiple wells for any container geometry, including microplates, petri dishes, and chambered slides.

*1.7 Organoid Fixation and Immunofluorescent Staining:* To preserve matrix structure, MAGIC matrices were pre-treated before fixation to hold bioprinted structures in place. Samples were first incubated for 20 min with 25 mM $CaCl_2$ in sterile milliQ $H_2O$ to "lock" the alginate microgels. Then, a layer of warm liquid 0.5% agarose was poured over the samples and allowed to cool for 5 minutes at room temperature followed by 5 minutes at 4 °C. Samples were then fixed in 2% PFA for 45 min at room temperature and washed with PBS-glycine 3x, 20 min each followed by washing with PBS 2x, 20 min each. Samples were left overnight in 25 mM $CaCl_2$ solution, as long-term storage in PBS led to formation of insoluble calcium phosphate, which occluded the samples. The next day, fixed organoids were permeabilized with 0.5% Triton X-100 for 15 min at room temperature and blocked with blocking buffer for 2 h at room temperature or overnight at 4 °C. If using mouse-origin primary antibodies, samples were also incubated overnight with 1:50 dilution mouse Fab fragment (Jackson ImmunoResearch #115-007-003). Samples were then incubated with primary antibody in blocking buffer for 24-48 h at 4 °C, rinsed in wash buffer 3 times for 1 h at room temperature, incubated with secondary antibody in blocking buffer overnight at 4 °C, and rinsed in wash buffer 3 times for 1 hr at room temperature. DAPI was added for 30 min and organoids were washed once more for 30 min in PBS. Finally, samples were cleared overnight in RapidClear 1.52 (SunJin Lab) at room temperature before imaging. Primary antibodies used include rat anti-ECCD2 (Thermo Fisher 131900), rabbit anti-LYZ (Thermo Fisher 129680), rabbit anti-CHGA (Novus Biologicals NB120-15160B), and chicken anti-GFP (Aves Labs GFP1010). Secondary antibodies were raised in goat and included Alexa Fluor 488 (Thermo Fisher A11039), Alex Fluor 568 (Thermo Fisher A11011), and Alexa Fluor 647 tags (Thermo Fisher A21247).

*1.8 Microscopy:* Live imaging of bioprinted structures during extrusion, which allowed for iterative human-in-the-loop adjustments to print parameters, was achieved using a Leica DMI8 programmed on the same custom GUI as the bioprinter components. After printing, live organoids were imaged using either a Zeiss Axio Observer Z1 with a Yokogawa spinning disk or GE Healthcare IN Cell Analyzer 2200 confocal microscope. Brightfield and live cell fluorescence (membrane tdTomato, green fluorescent protein) were captured using 5x/NA 0.25 or 10x/NA 0.3 air objectives in controlled environmental chambers held at 37 °C and 5% $CO_2$. Fixed organoids were imaged using a Zeiss LSM800 confocal microscope equipped with 20x/NA 0.8 LD air or 25x/NA 0.8 multi-immersion objectives. Image acquisition and stitching was controlled using ZEN 2.3 (2011) software. Subsequent image analysis, including background subtraction, filters, and thresholding was performed using custom macros in Fiji/ImageJ. Any image alterations, such as brightness & contrast adjustments or thresholding, were kept consistent across conditions in a given experiment for analysis.

*1.9 Organoid Size and Crypt Analysis*: Quantification of organoid area and crypt number were made based on binary masks constructed using Fiji/ImageJ. Masks were achieved by taking maximum z-projections of all imaged organoids in the tdTomato+ channel, adjusting contrast, applying a Gaussian blur (5-20 μm), and adaptively thresholding. Following conversion to a mask, holes were filled and organoids were identified with Analyze Particles (excluding objects < 5000 $\mu m^2$, parameter=exclude add). To quantify organoid area, the area metric was extracted with ROI Manager (measure). To further quantify the number of crypts, we employed a MATLAB crypt counting software developed by Montes-Olivas et al[11]. Here, had to filter out smaller manually seeded organoids to maintain crypt-counting accuracy. Consequently, the organoids included in this analysis were biased toward larger organoids, of similar size to the bioprinted condition.

For microwell comparison experiments, Caco-2 cells were prepared as described above and counted by hemocytometer to calculate the desired cells seeded per EZSphere (AG4860-900SP) microwell. Cells were centrifuged to pellet at the base of the microwells before Matrigel was added on top of the wells and allowed to solidify. Media was then added to the wells and seeded spheroids were imaged and analyzed as described above. For area quantification, the median 80 detected objects per well were used from 2 wells per seeding condition to match the 80 microwells/well EZSphere density and to exclude imaging artifacts.

*1.10 Human umbilical vein endothelial cell (HUVEC) culture and matrix preparation for bioprinting:* HUVECs (Lonza) were cultured in EGM-2 and used between passages 4 and 6 as described previously[12]. mCherry-HUVECs were created by transducing cells with a pSicoR-EF1a-mCherry lentivirus. All lentiviruses were made by the UCSF Viracore. Transduced cells were sorted on a BD Aria II flow cytometer. For bioprinting experiments, confluent HUVECs were digested using TrypLE Express (Gibco) for 10 min at 37 °C. Cells were washed in D-PBS with 2 mM EDTA and filtered using a 40 μm cell strainer. Single-cell solutions were kept on ice until printing. MAGIC matrices with 0.5 wt% alginate mixed at a 1:1 added volume ratio to Matrigel were further mixed with a neutralized stock solution of 8.5 mg/mL rat tail collagen I (Advanced Biomatrix) to achieve 1 mg/mL collagen in MAGIC matrix. This matrix was used for printing at 4 °C before ECM cross-linking at 37 °C.

*1.11 Bioprinted Tube Perfusion:* Bioprinted tubes were allowed to self-organize and form lumens for 3–7 days, with media changes every other day. Gut tubes were generally printed into #1.5 coverglass-bottomed chambered slides with one chamber, to allow access with a glass capillary and micromanipulator. Aluminosilicate glass micropipettes with a long tether were prepared using a P-97 micropipette puller (Sutter Instruments). The pulled pipettes were cut 3–5 mm from the tip to get 10–25 mm-diameter pipettes with jagged ends and filled with PBS. Once patent lumens were visible, one end of a tube was cut with a razorblade to create an opening, and the other end was pierced with the capillary mounted on a Narishige MM-89 micromanipulator connected to a syringe. Applying pressure to the syringe induced liquid and debris flow toward the open end of the tube. Images were acquired using a 10x/NA 0.25 air objective on a Zeiss Axiovert 200M running SlideBook software. Tube diameter as a function of time was measured by manual thresholding of the tube and dividing thresholded area by imaged tube length to get an average tube diameter. This was performed over each frame of a video over multiple pulse cycles.

*1.12 Human primary mammary organoid bioprinting and analysis:* Deidentified normal, finite lifespan primary human mammary epithelial cells (HMEC) were provided by Drs. Martha Stampfer and James Garbe (Lawrence Berkeley National Laboratory). HMEC were cultured from tissues removed during reduction mammoplasties, and expanded to fourth passage in M87A medium as described previously[13]. HMEC from tissue donated by a 19 year-old individual with bilateral breast hypertrophy (240L) were used for all experiments. All HMEC were cultured in complete M87A medium with Penicillin-Streptomycin (100 U/mL) at 37 °C with $CO^2$ up to 80-90% confluency. Fourth passage HMEC were transduced with lentivirus upon thawing at a multiplicity of infection of 13 (to target 40-60% transduction efficiency) into a half volume of M87A medium containing 2 µg/ml polybrene (Millipore-Sigma #TR1003). After 3 hours, M87A medium was added to full volume. After 24-48h, the virus-containing medium was discarded and replaced with fresh M87A medium. Cells were grown up to 80-90% confluency (5-7 days). Transduced cells were isolated by FACS based on GFP or mCherry expression. To obtain a printable single-cell solution, 80-90% confluent HMEC were digested using TrypLE Express (Gibco) for 10 min at 37 °C. Cells were washed with D-PBS with 2 mM EDTA and filtered using a 40 µm cell strainer. GFP- or mCherry-expressing HMEC were sorted into myoepithelial and luminal populations and recombined into bioinks using 1:1 and 2:1 LEP:MEP or all-MEP compositions. Bioprinted HMEC organoids were analyzed for sorting one day after printing using ilastik[14] to segment based on fluorescence and custom Fiji scripts to assess boundary occupancy.

*1.13 Human iPSC culture, cortical organoid differentiation, dissociation, and cell proportion analysis:* iPSCs from three different donors were used (28126, 20916B, 13234)[15,16] were maintained on Matrigel-coated plates and cultured in StemFlex (Thermo Fisher #A3349401). To passage, cells were lifted with PBS without calcium or magnesium supplemented with 0.5 mM EDTA. For differentiation, iPSCs were lifted using PBS-EDTA and resuspended in Neural Induction Media containing GMEM (Gibco #11710035), 10% Knockout Serum Replacement (Gibco #10828028), NEAA diluted 1:100 (Gibco #11140050), Sodium Pyruvate diluted 1:100 (Gibco #11360070), 5 mM 2-mercaptoethanol (Sigma Aldrich #M6250), and 100 µg/mL Primocin supplemented with 5 µM SB431542 (Tocris #1614), 100nM LDN-193189 (Sigma Aldrich #SML0559), and 3 µM IWR1-endo (Cayman Chemicals #13659). Cells in Neural induction media were moved to 6-well low attachment plates (Corning #3471), with a media change on day 3 with CEPT, and day 6 without CEPT. From day 9-25, organoids were cultured in Maintenance Media 1: 50% DMEM/F12 with Glutamax (Gibco #10565042) and 50% Neurobasal (Gibco #21103049) with B27 without vitamin A (Gibco #12587001), N2 (Gibco #17502048), NEAA diluted 1:100, Glutamax diluted 1:200 (Gibco #35050061), and 55 µM 2-mercaptoethanol supplemented with 10 ng/mL each FGF (Peprotech #100-18B) and EGF (Peprotech #100-47). Media was changed every 2-3 days. From days 26-35, media was changed without FGF and EGF. From day 35 onward, organoids were cultured in Maintenance Media 2: Maintenance Media 1 supplemented with B27 with vitamin A (Gibco #17504001), instead of B27 without vitamin A. For printing, organoids were dissociated on D49 using 20 units of Papain (Worthington #LK003178) with 5% Trehalose in HBSS for 30 minutes at 37 °C. DNase was added, and organoids were incubated for another 15 minutes at 37 °C. Papain was quenched using Albumin-ovomucoid inhibitor (Worthington # LK003182) and cells were filtered through a 40 µm cell strainer. Cells were spun down, resuspended, and counted.

For cell proportion quantification, 20 µM optical sections were used with maximum Z projection and LUT adjustment consistent for all images. CellProfiler[17] version 4.2.4 was used to quantify the number of cells positive for each protein of interest. Briefly, IdentifyPrimaryObjects was used to identify cells positive in each channel. Then RelateObjects and FilterObjects were used to assign a parent-child relationship between DAPI and each additional channel, removing cells with positive signal in one channel but not for DAPI. Percentage of cells positive for each marker were then calculated using these metrics.

*1.14 Triple-negative breast cancer patient-derived organoid transduction & transfection:* TORG139 patient-derived triple-negative breast cancer (TNBC) organoids were generated and cultured as previously described in type 1 mammary organoid medium[18–20]. To obtain a printable single-cell solution, organoids were digested using TrypLE Express (Gibco) for 10 min at 37 °C. Cells were washed in D-PBS with 2 mM EDTA and filtered using a 40 µm cell strainer. Single-cell solutions were kept on ice until printing. For transduction experiments, TNBC organoids were mechanically passaged and either incubated in suspension for 2 hours in media containing GFP-expressing lentivirus or replated in Matrigel or MAGIC matrix before adding media containing GFP-expressing lentivirus with a multiplicity of infection of 5 ($10^6$ viral particles per sample) and 2 µg/mL polybrene (Millipore-Sigma #TR1003). Bioprinted TNBC organoids were transduced in the same conditions with virus-containing media left on top of the matrix in a 96 well plate. Viral media was discarded 2 h (suspension) or 16 h (matrix-embedded and bioprinted) after transduction. Organoids were imaged at day 5 post-transduction and GFP expression was quantified by segmenting individual organoids and sub-segmenting those with GFP signal using ilastik[14]. Following manufacturer's instructions, bioprinted TNBC organoid arrays were transfected using Lipofectamine RNAiMAX (Thermo Fisher) in Type 2 medium[21] with 1% FBS and with 5 or 10 pmol of a Cy3-conjugated 36mer single-stranded non-coding small RNA (MW = 11.9 kDa) for 24, 48, or 72 h starting 3 days after printing. Images were taken with an Echo Resolve microscope.

*1.15 Gamma-Secretase Inhibition Experiment and Analysis: Atoh1$^{CreERT2}$:Rosa26$^{tdTomato}$* gut organoids were isolated from proximal mouse small intestine and cultured as described above. Organoids were bioprinted or manually seeded and immediately cultured in ENR supplemented with either 50 µm *N*-[*N*-(3,5-Difluorophenacetyl-L-alanyl)]-(*S*)-phenylglycine *t*-butyl ester (DAPT, +treatment) or a DMSO vehicle control (–treatment). The media was changed on the second day, and 4-hydroxytamoxifen (4-OHT, Sigma-Aldrich H7904). On the third day after seeding, organoids were imaged using equivalent settings in high-throughput on a GE Healthcare IN Cell Analyzer 2200 Confocal Microscope in brightfield and 568 nm channels equipped with a 10x/NA 0.35 air objective. For microscopy images, a max intensity projection was created in the 568 nm channel to show tdTomato+ signal, and a representative central focal plane was chosen in brightfield. For analysis, a custom Fiji macro was generated to identify and segment individual organoids. The tdTomato+ volume of each individual organoid was then calculated in Fiji using a custom thresholding macro. Bootstrapping and subsequent p-value quantification between treated and untreated conditions for either printing or manual seeding was performed using a custom R script[22] with 512 iterations at each number of pairwise organoids.

*1.16 Statistical Analysis:* Sample numbers for a given experiment are provided in each figure legend and were always $n \geq 3$ independent replicates. Statistical analyses included non-parametric t-tests, one-way ANOVAs with Tukey's or Dunnett's multiple comparisons, normality tests, conventional power analysis using $\alpha = 0.05$ and $\beta = 0.2$, and bootstrapping as described above. Statistical analysis was performed using GraphPad Prism 10 (t-tests, ANOVAs, normality, power analysis) or R (bootstrapping).
